# The impacts of climate variability on the niche concept and distributions of species

**DOI:** 10.1101/2024.10.30.621023

**Authors:** Emilio Berti, Ángel Luis Robles Fernández, Benjamin Rosenbaum, Townsend A. Peterson, Jorge Soberón, Daniel C. Reuman

## Abstract

Ecological demographers know that year-to-year climate variability influences the long-term growth of populations and thus their viability. Despite this, species distribution models (SDMs), widely used to project species’ geographic distributions based on climate, typically ignore inter-annual climate variability. Here, we show that climate variability plays a crucial role in determining current and future distributions of species. We achieved this by developing a new SDM framework, XSDM, that accounts for variability when assessing the ecological niche and distribution of species. XSDM outperforms traditional SDMs in simulation studies. Using XSDM, we assessed the impacts of variability on 10 example species. Variability reduces species potential distributions by an average of 22%, up to 45%. Moreover, sensitivities of distributions to potential changes in average temperature and in its variability were comparable in magnitude. To avoid biases, future SDMs should consider the effects of variability through a demographic approach such as XSDM. As global change alters climate variability, e.g., through increased frequency of extreme events, XSDM provides better tools for countering biodiversity losses.

## 3 Main

One of the main consequences of climate change is the modification of the distributional ranges of species, as species move to track altered conditions [1]. Redistribution of biodiversity has major conservation consequences, and can negatively affect ecosystem services, such as productivity and carbon storage, that are essential to human societies [2]. Therefore, effective efforts to determine how species geographic distributions will respond to climate change are urgently needed. To assist with this challenge, a variety of ecological niche modeling (ENM) and niche-based species distribution modeling (SDM) approaches are commonly applied [3]. ENMs and SDMs relate environmental conditions (mostly climate) to observation records of a species, and have been used to estimate the current and future distributions of species [4]. However, a major limitation of current ENMs/SDMs is they do not account for inter-annual climate variability, even though variability is known by demographers to play a crucial role in population dynamics and therefore possibly also in species distributions [5, 6, 7, 8, 9]. Climate change is also altering climate variability, e.g. by modifying the intensity, frequency, and duration of extreme events such as heat waves and floods [10, 11]. Addressing the shortcomings of current SDMs with respect to climate variability therefore seems important for better understanding the impacts of climate change on biodiversity.

Inter-annual climate variability affects population dynamics by inducing variability in annual net rates of population growth [12]. Importantly, the effects of climate variability on a population can be quantified using stochastic demography, a field of research which examines how the performance of a population is influenced by fluctuations in population vital rates [13]. A key concept of stochastic demography is the long-term stochastic growth rate (ltsgr), which helps describe the long-term viability of a population [13], with larger values generally indicating a population which is more likely to persist in a locale. For example, for a population following the simple Lewontin-Cohen model [14],

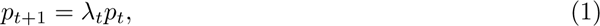

where *t* is the year, *p_t_* the number of individuals of the population in year *t*, and *λ_t_* the net growth rate in year *t*, the ltsgr is, approximately [15],

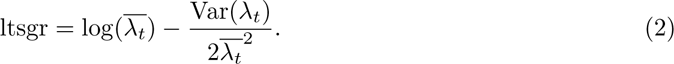

Here, 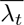 is the average growth rate and Var(*λ_t_*) is the variance of growth rate across the years. We use the natural logarithm here and throughout the paper. Eq. 2 illustrates how inter-annual climate variability can influence long-term population viability: higher climate variability induces elevated inter-annual variance of growth rate, Var(*λ_t_*), thereby lowering the ltsgr. See Figs 1 and 2 for further elaboration of this example.

**Figure 1:**
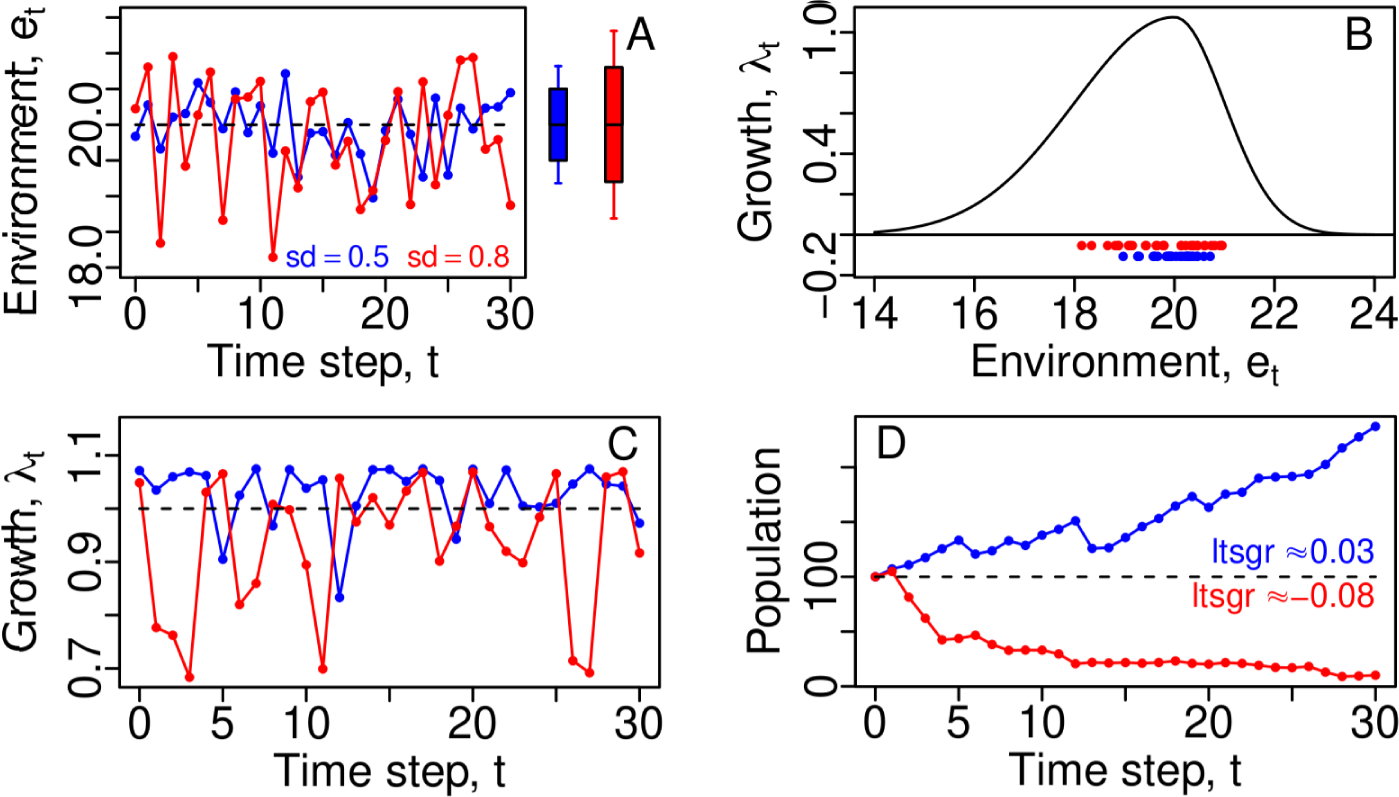
Modelbased illustration of how interannual climate variability can affect population viability. Two distinct locations (red and blue points/lines on all panels) are assumed to have identical average climate conditions, *e̅_t_*, but different degrees of interannual variability (A; sd = standard deviation). If modeled populations in each location follow the Lewontin-Cohen model (see text) and the annual net growth rate, *λ_t_*, of each population is a function of local climate as in B, then differences in climate variability between the locations can produce large differences in time series of growth rates (C), which lead to dramatic differences in population survival outcomes (D). The sign of the longterm stochastic growth rate (ltsgr; see text) accurately reflects population viability in these two scenarios (D). The function in B is given by eq. (5) in Methods.

**Figure 2:**
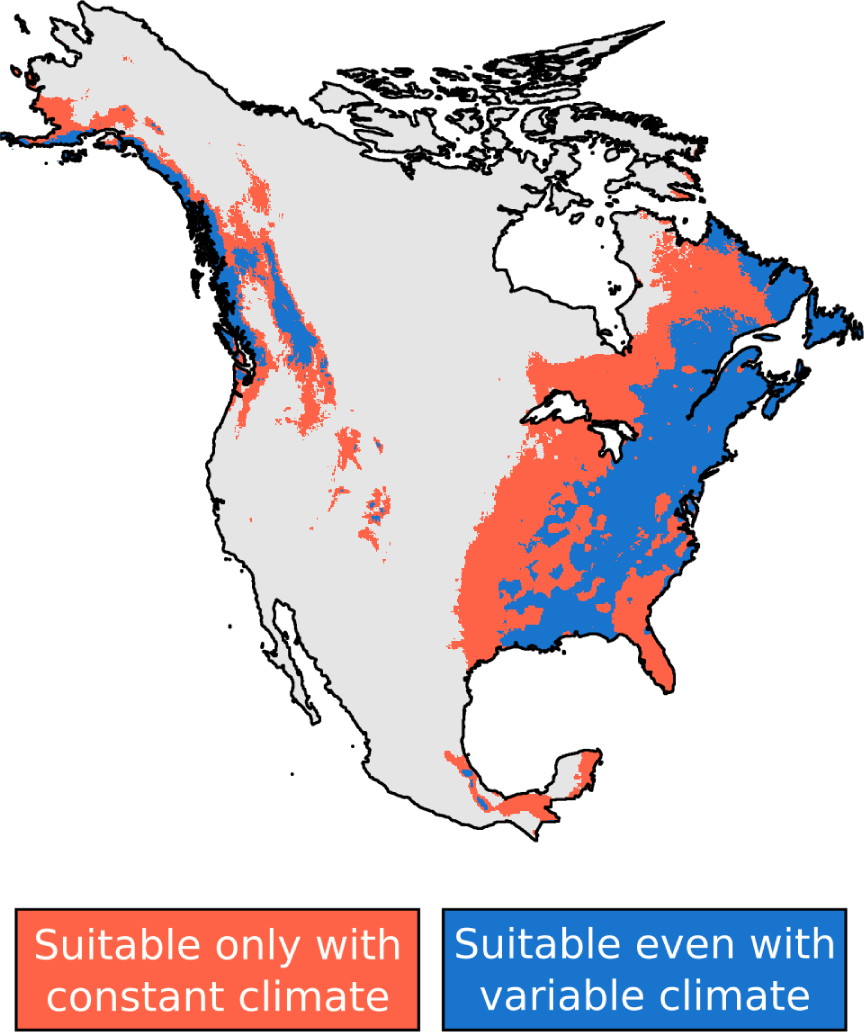
Modelbased illustration of how interannual climate variability can drastically affect species’ geographic range. Modeled populations in each location again follow the Lewontin-Cohen model (see text and Fig. 1), with the annual net growth rate, *λ_t_*, in a location given as a function of *e_t_*, with *e_t_* being the precipitation of the driest quarter in the location in year *t* (the specific function used is again eq. (5) in Methods). When accounting for variability, the species geographic range can be considered to be locations (blue) for which the longterm stochastic growth rate (ltsgr; see text) is greater than zero. When not accounting for variability the range consists of the locations (combined red and blue, so that red shows the reduction in range due to variability) for which the growth rate, *λ_t_*, at the average climatic condition is greater than 1, i.e., *e̅_t_*. Traditional SDMs typically use average climate to predict species geographic range. Parameters of eq. (5) used for this example were *µ* = 3000mm, *σ_L_* = 200mm, *σ_R_* = 2 · 10*^7^*mm, and *λ_max_* = 1.1.

Though such ideas are fundamental to stochastic demography, current ENMs/SDMs usually rely on long-term climatic averages, which neglect inter-annual climate variability (equivalent to setting 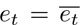 in the Fig. 1 example). Traditional ideas of the fundamental niche underlying common SDMs have also commonly been based on deterministic growth, e.g., the *n*-dimensional hypervolume of Hutchinson [16] is commonly operationalized by requiring that average environmental conditions be suitable for a deterministic population growth rate to be greater than 1 [17]. Whereas some SDM efforts account for intra-annual climate variability, e.g. using bioclimatic variables linked to seasonality, such variability is distinct from the inter-annual variability that we consider: annual net growth rate of a population may depend on intra-annual variability, but the ltsgr is influenced by inter-annual variability in annual net growth. Additionally, the life histories of a great many organisms are evolved to cope with seasonal variation, while populations remain sensitive to inter-annual variation in important seasonal variables such as winter temperature or spring rainfall [18].

Here, we show that inter-annual climate variability substantially impacts the potential distributions of 10 North-American species chosen as examples. And we show that sensitivities of species’ distributions to changes in average temperature and to changes in temperature variability are comparable in magnitude, so that the latter should not be ignored when projecting future species distributions. We arrive at these conclusions by developing a novel framework, called XSDM, that integrates concepts of stochastic demography into SDMs. XSDM does not require extensive demographic information, is computationally efficient, relies on easily accessible data, and shows better performance in simulation studies than traditional SDMs. Overall, our results show that including inter-annual climate variability is key to more effectively conceptualizing the niche of species and more accurately understanding their current and future distributions.

### 4 XSDM: a novel demographic SDM to account for climate variability

The XSDM approach infers the ecological niche of a species from occurrence data and time series of climatic conditions, thereby obtaining the species’ potential geographic distribution. Inspired by important previous work [19, 20, 21], XSDM contains two modules (Fig. 3): 1) a demographic response model, where a *growth-environment function* describes how inter-annual variation in climate translates into variation in population net growth rate – this part of XSDM is used to compute the local ltsgr; and 2) a habitat suitability model, where a *suitability-link function* is assumed to relate the local ltsgr to the suitability of the local environment for the species. Crucially, XSDM directly quantifies the effects of climate variability on a species’ potential distribution through the use of the ltsgr. During model fitting, parameters of both modules are jointly adjusted to maximize agreement between model-predicted habitat suitability and actual species occurrence records. See Methods for additional details of how XSDM works.

**Figure 3:**
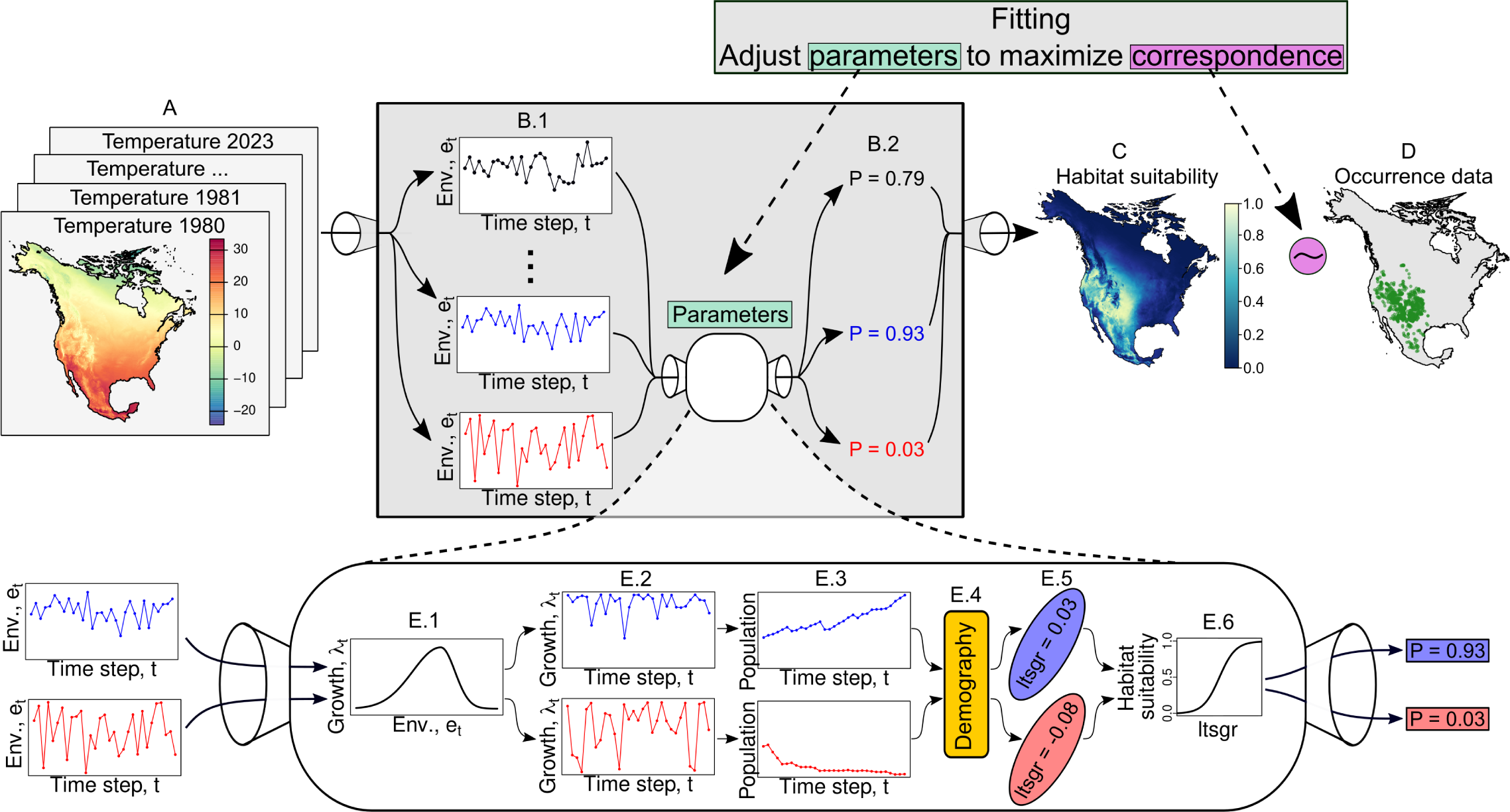
Conceptual illustration of how XSDM works. XSDM takes as input maps of annual values of one or more climate variables (A). It then breaks these into separate time series for each location (B.1) and passes them to the demographic response model, specifically to the growth-environment function (E.1), which describes the influence of environment (*e_t_*) on annual net growth rate (*λ_t_*) and has adjustable parameters. This generates time series of growth rates (E.2), which are related to population growth trajectories (E.3). Demographic theory (E.4) is used to calculate the ltsgr (E.5), which incorporates the influence of climate variability. Larger values of ltsgr correspond to better growth conditions and imply higher habitat suitability, modeled through a suitability-link function with adjustable parameters (E.6). Habitat suitability values (B.2) are aggregated across all locations to produce a suitability map (C). The demographic and habitat suitability models are fitted jointly in XSDM, adjusting the parameters of the growthenvironment and the suitabilitylink functions (E.1 and E.6) in order to maximize the match between predicted suitability (C) and occurrence data (D). This figure, for pictorial simplicity, assumes throughout that one climate variable only, *e_t_*, controls *λ_t_*; but multiple variables are typically used.

We performed two extensive simulation studies (Methods) with virtual species to assess to what extent XSDM could be used to accurately infer parameters, and to compare how XSDM performs relative to existing, commonly used SDMs. Model fitting of XSDM accurately recovered important model parameters (Extended Data Fig. 1) and the suitability for virtual species of each simulated habitat patch (Fig. 4, A; Extended Data Fig. 2), with greater accuracy for larger sample sizes, as expected (Fig. 4B). Moreover, XSDM outperformed other commonly used SDMs (Fig. 4, C; Extended Data Fig. 3), producing results that were both more accurate and less uncertain.

**Figure 4:**
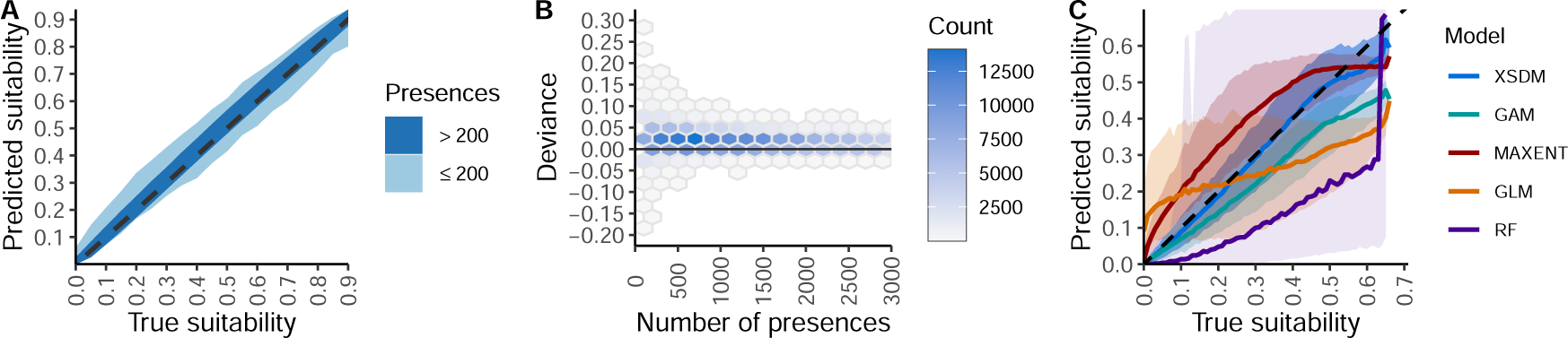
Assessments of XSDM’s ability to infer true simulated habitat suitability (A, B), and performance of XSDM compared to other SDMs (C). (A) XSDM accurately predicts the true climatic suitability of habitat patches for virtual species. Shades show the 95% quantile range of predicted suitability values. The dashed black line is the 1:1 line indicating perfect predictions. Colors show two levels of sample size (numbers of positive detection records of the focal species), highlighting that XSDM performs very well with a number of presences ≥ 200 but can also perform well for fewer presences. See Extended Data Fig. 1 for recovery-accuracy results for particular model parameters. (B) “Deviance” in XSDM predictions, i.e. the difference between predicted and true habitat suitability, decreases with increasing sample size, as expected. The intensity of blue color shows the number of virtual habitat patches from simulations in each hexagon. (C) XSDM outperforms commonly used SDMs, namely generalized additive models (GAM), Maxent, generalized linear models (GLM), and random forests (RF). See Methods, Extended Data Figs 2 and 3, and Supplementary Information for details.

### 5 Climate variability constrains the distributions of species

To highlight how climate variability can affect species’ distributions and illustrate the possible applications of XSDM, we modeled the potential distribution of 10 species (2 birds, 1 insect, 3 mammals, 3 plants, and 1 reptile) chosen from checklists of North America species (Methods). On average, inter-annual climate variability reduced the potential distributions of species by 22% (median = 20%). The strongest effect was for the reptile *Ophisaurus ventralis*, for which the distribution was reduced by 45% (Fig. 5). Of the species we examined, the bird *Nucifraga columbiana* was the least affected by climate variability, with its potential distribution barely reduced at all (1%).

**Figure 5:**
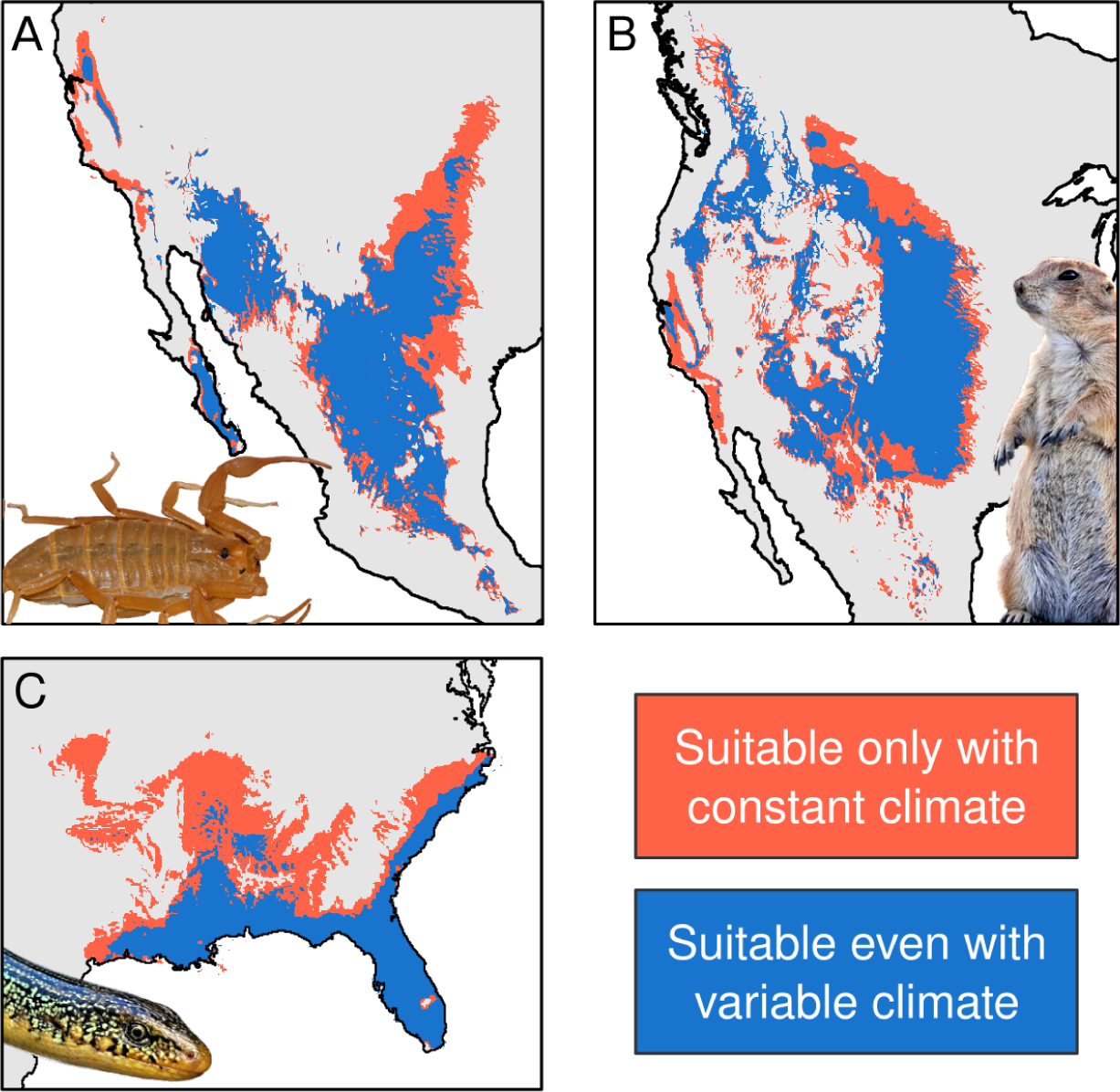
Potential distributions, as projected by XSDM, for (A) the Arizona bark scorpion (*Centruroides sculpturatus*); (B) the black-tailed prairie dog (*Cynomys ludovicianus*); and (C) the Eastern glass lizard (*Ophisaurus ventralis*). Colors show the potential species distribution when accounting for interannual climate variability (blue) and when neglecting it (the union of the blue and red, so that red shows the reduction in range due to variability). Projections show areas of suitable climate and do not account for possible dispersal barriers or range limitations which may be imposed by other species with which the focal species interacts, as commonly done also in traditional SDMs (but see Discussion for potential improvements). Original animal photos by Andrew Meeds, Cephas, and Ignioapathy (CC BYSA 4.0).

### 6 Species distributions are likely sensitive to future changes in variability

Species potential geographic ranges were comparably sensitive to increases in inter-annual variability of temperature and to increases in average temperature (Fig. 6), suggesting that future species ranges may be altered in two ways by climate change, both similarly important but only one accounted for by typical SDMs. Sensitivities of geographic ranges to increases in average temperature were positive for seven of our ten species (Fig. 6), which thus were projected to gain distributional range as a result of small increases in average temperature (Extended Data Figs 4, 5). Sensitivities were negative for three species, which thus were projected to lose range; sensitivity was particularly high in magnitude for *O. ventralis*. Sensitivities to increases in inter-annual variability of temperature were negative for all species and had particularly high magnitude for *O. ventralis*, which was projected to lose around 75% of its current range for an increase of inter-annual standard deviation of temperature of one degree Celsius (Fig. 6). Four species were more than 2 times as sensitive to an increase in average temperature as to an increase in inter-annual variability. Two species were more then 2 times as sensitive to an increase in inter-annual variability as to an increase in average temperature. For the four remaining species the sensitivity to average temperature was comparable in magnitude to the sensitivity inter-annual variability, namely between 1*/*2 and 2 times. Sensitivities reported here measure percent change in the potential geographic range of a species per degree Celsius change in average or standard deviation of temperature (Methods).

**Figure 6:**
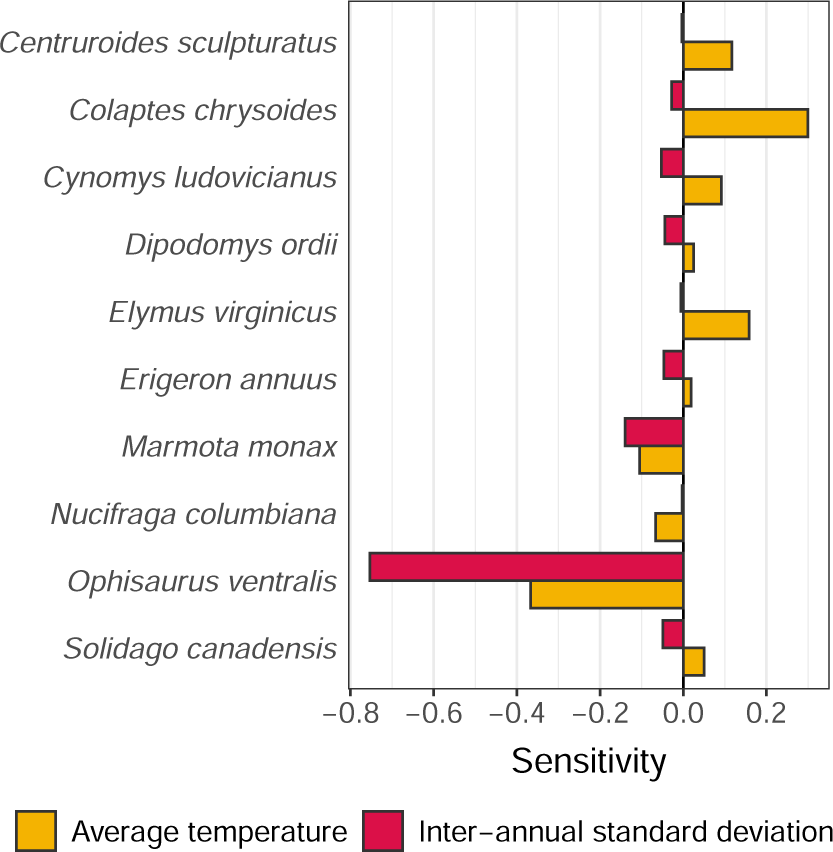
Sensitivities of species geographic ranges to increases in average temperature or interannual standard deviation of temperature for the 10 species studied. Negative sensitivities indicate percentage range loss projected to result from a small increase in average or standard deviation of temperature; positive sensitivities indicate a corresponding increase in range. Sensitivities are in units of *^◦^*C*^−1^*. Numeric results are in Supplementary Table 1.

## 7 Discussion

We showed that inter-annual climate variability can strongly influence species’ distributional patterns, though this influence has been largely ignored by standard SDMs. We proposed the novel framework XSDM to help resolve this limitation of SDMs and to begin to quantify the current and future influence of climate variability on distributions. Key to these advances was the integration of theory from stochastic demography into niche and distribution modeling. Because XSDM takes into account stochastically fluctuating environments, our framework moves away from the deterministic view of the fundamental niche which has been common in past work, i.e., where the niche has been considered to comprise values of the average environment for which a deterministic growth rate indicates population growth. Instead, our approach focuses on the sets of stochastic environments for which the long-term stochastic growth rate is positive [19], and the locations in geographic space exhibiting such stochastic environments.

Several important studies have advocated for process-explicit SDMs based on demographic theory, and have helped inspire our work [12, 19, 22, 23, 20, 24, 25]. Keith et al. [26] and Eckhart et al. [27] showed that demographic approaches to SDMs can be constructed, and similar approaches were later used by others [28, 29]. But those models were both limited in their applicability to very extensively studied species and used deterministic demographic approaches that did not account for climate variability. Perez-Navarro et al. [6] included inter-annual variability in their model, showing that it likely plays a role in the distribution of *Pinus halepensis*. But their approach did not model demographic processes. Importantly, Pagel and Schurr [20] and Schurr et al. [21] proposed a research agenda integrating population processes into SDMs. Their framework has been very valuable, but is so data and computationally hungry as to be difficult to apply. Efficiency gains of XSDM relative to the method of [20] are due to the use of tools from stochastic demography. For instance, a typical time it took us to fit a single XSDM model with GBIF data for one of our example species was about 20-25 minutes on a compute node with 54 cores and 16 Gb RAM. It should be straightforward to run XSDM for numerous species on a typical computing cluster. Additionally, a body of literature has tried to relate environmental suitability evaluated through ENMs with species’ population density, with contrasting results. Theory predicts a positive relationship between environmental suitability and population density, confirmed in some studies [30, 31, 32] but not others [33, 34] (see [35] for caveats). Our approach could be expanded, in future work, to relate environmental suitability as estimated through stochastic demography to population density, e.g. by relating the ltsgr to density (Methods). This may help reconcile contrasting evidence from previous studies.

Karger et al. [36] included inter-annual variances of environmental variables as predictors in traditional SDMs. They showed that model performance was thereby only slightly improved, creating an apparent contrast with the large importance of climate variability we demonstrated. We interpret this contrast as arising because the models of [36] were traditional, non-process-based SDMs. Variance was used by their models as just another predictor which could be used to explain species’ suitability. However, the influence of climate variability on population viability is known to be complex, having been quantified through much research in stochastic demography [13, 15]. It may not be possible to adequately account for this influence with the simple phenomenological models of standard SDMs, even when environmental variability data are provided to those models.

XSDM shares some of the same assumptions as traditional ENMs/SDMs. XSDM assumes that observed occurrences of the modeled species are limited most prominently by environmental constraints, not by dispersal limitations or by biotic interactions such as competition or predation from other species [17, 19]. We help mitigate this assumption, in a manner also often used by other ENMs/SDMs, by selecting pseudoabsences only within 100km of the convex polygon containing presence records, to increase the probability that pseudoabsences occur in locations which are environmentally unsuitable instead of merely inaccessible. Projections of XSDM are projections of potential or environmental habitat suitability, and say nothing about whether the focal species could successfully disperse to a potential location, or whether it may be prevented by biotic interactions from establishing [4, 37]. Like most other ENMs/SDMs, XSDM does not account for source-sink or other metapopulation dynamics. It is expected that some of the efforts which have been made to ameliorate or circumvent these assumptions for traditional ENMs/SDMs could also be applied to XSDM.

Several potential improvements to XSDM seem worth considering in future work, including the following. First, the (st)age-structured demography of a species could be explicitly modeled using stochastic matrix models [13], allowing for different environmental sensitivities for different (st)ages. To simplify the presentation of main ideas, we here considered the Lewontin-Cohen model to be an adequate starting point; future studies could build technical advances on top of the conceptual foundations we here established by extending XSDM to stochastic matrix models. Second, the “infinitesimal variance” and temporal autocorrelations of population vital rates, both known to demographers to influence population viability [13, 38], could straightforwardly be incorporated into XSDM. Third, it may be possible to integrate into XSDM the potential for a species to exhibit local demographic adaptations. For example, the optimum climate, *µ* (eq. 5), for a species could be modeled to change gradually along a geographic gradient. Fourth, additional factors that do not relate to climate could be allowed to influence the suitability-link function, e.g. habitat type and human land use. The simple version of XSDM used here was sufficient for an initial investigation of the importance of variability to species distributions, but extensions such as those listed above may be beneficial for some future applications. Importantly, the XSDM framework allows for future researchers to test whether each extension is statistically warranted.

The fundamental niche, *N_F_*, of a species has been defined [16, 39, 40, 17] as the set of all points in an *n*-dimensional environment space for which a species’ population growth rate is positive. Specifically, in the language used by Sobeŕon and Peterson [39, 40, 4], this is the Grinnellian fundamental niche and the environmental variables are taken to be scenopoetic [37]. One useful lesson emerging from our results is that the *n*-dimensional environment space ought to also include axes relating to inter-annual environmental variation through time, as well as the traditional axes representing mean values of environmental variables; but we think a slightly more abstract conceptualization will be even more useful for guiding future research. Consider, instead, the *n*-dimensional environment space to be the space of all *p × T* matrices representing time series of the *p* relevant environmental variables over a period of *T* years; so *n* = *pT*. Thus, points in environment space are entire temporal trajectories of environmental variables over an appropriate time period for the species (Fig. 3, E.2). This viewpoint is similar in some ways to those of [9, 40], though, importantly, the viewpoints of [40] are intra-annual and ours are inter-annual. The species growth rate we then use to delineate *N_F_* is the ltsgr (Fig. 3, E.5). Our *pT* -dimensional space projects onto a smaller, 2*p*-dimensional space representing the means (the climatological normals commonly used in SDMs) and variances (as used, for example, by Karger et al. [36]) of all *p* environmental variables, so includes all the information that the smaller space includes. But the larger space also retains information about, for instance, temporal autocorrelations, known to demographers to be important for population viability (see above). Compressing the environment space from a *pT* -dimensional space to a smaller 2*p*-dimensional space throws away information that may help quantify species performance under actual conditions. This is another viewpoint on the explanation, above, of why the models of Karger et al. [36] only showed slight performance improvements with the inclusion of variance information, whereas our models demonstrated the substantial importance of variability for species’ range.

Understanding when and how climate variability influences the distribution of species is important not only for fundamental scientific advancement, but also for helping mitigate the negative effects of climate change. Climate change includes alterations to the frequency and intensity of extreme events [10, 11]; and these are aspects of climate variability. Our results indicate that species geographic ranges may well often be at least as sensitive to changes in the inter-annual variability of temperature as to changes in average temperature. Therefore, climate change may strongly alter species distributions not only through its effects on average temperature [1], but also through its effects on extreme events and other aspects of climate variability. Prior work indicates that changes in climate variability may cause greater changes in species ltsgrs than do changes in climate means [41, 42]. Through XSDM, we here developed the foundations for an improved understanding of these two distinct pathways by which climate change can alter species distributions.

## 8 Methods

### XSDM dynamical model

Populations were modeled in a location by the Lewontin-Cohen model [14],

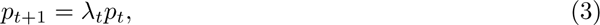

where *p_t_* is the population at time *t*, and *λ_t_* is the net population growth rate at time *t*, which depends on aspects of the environment at the location at time *t*. This modeling choice effectively assumes that, for the purposes of determining baseline environmental suitability of a location, density dependence can be neglected; similar to an assumption that a habitat is suitable if a population can increase there when rare. This is analogous to the invasion criterion used in invasion biology [43] and to other similar assumptions commonly used for conservation applications of stochastic demography [13, 44, 45]. See Discussion for future extensions of XSDM that could incorporate population (st)age structure.

The viability of the population (Fig. 3, E.4-5) is assessed by the long-term stochastic growth rate (ltsgr), which is

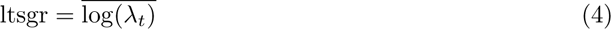

for the model of eq. 3 [13]. The overbar denotes a mean over time. The ltsgr is well known to relate closely to population extinction risk [38] and is widely used in quantitative conservation biology [44]; see introductory texts for additional background on the ltsgr [13, 44, 45]. The approximation of eq. 2 was used in the Introduction to clarify concepts, but eq. 4 was used elsewhere.

### XSDM demographic response model

The growth-environment function (Fig. 1, B; Fig. 3, E.1), describing how *λ_t_*is influenced by environmental factors, will typically depend on several environmental variables (e.g., *λ_t_* may depend on both spring temperature and rainfall). To simplify the presentation of main ideas, the univariate case only is described here, with the multivariate case described in Supplementary Information. For our univariate growth-environment function, we used an “asymmetric bell-shaped” curve with niche parameters *θ_1_*= {*µ, σ_L_, σ_R_, λ_max_*}, i.e.,

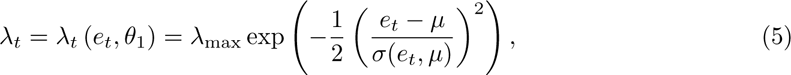

where *µ* is the optimal climatic condition for growth and *σ*(*e_t_, µ*) determines the shape of the curve via *σ*(*e_t_, µ*) = *σ_L_* if *e_t_* ≤ *µ*, and *σ*(*e_t_, µ*) = *σ_R_* if *e_t_ > µ*. This function can produce a large variety of shapes, e.g. symmetric (*σ_L_* = *σ_R_*) and asymmetric (*σ_L_* /= *σ_R_*) bell shapes as well as saturating (*σ_R_ >>* 1) responses (Supplementary Fig. 1).

### XSDM habitat suitability model

To relate the ltsgr to habitat suitability for the species, we used a sigmoid suitability-link function (Fig. 3, E.6) with parameters *θ_2_* = {*b, c, p_d_*}, i.e.,

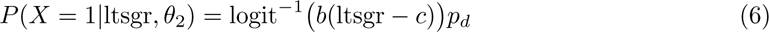

where logit*^−^*^1^(*x*) = 1*/*(1+exp(−*x*)), *p_d_* ∈ (0, 1) controls the maximum possible suitability, *c* controls the location of the center of the sigmoid, and *b* controls its sensitivity to the ltsgr.

### Identifiability

XSDM is “structurally non-identifiable” [46] when parameterized by {*µ, σ_L_, σ_R_, λ_max_, b, c, p_d_*}, i.e., these parameters cannot all simultaneously be determined by fitting. Parameter-space dimension was reduced in a manner that addressed this problem and permitted practical application of the model. The reduced space involved the parameters 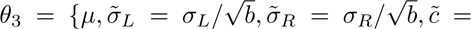 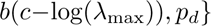. The re-parameterized model was both structurally identifiable and “practically identifiable” [46] for our datasets. Details in Supplementary Information.

### Model fitting

A likelihood function was formulated based on the model specifications above, and the parameters *θ_3_* were then jointly estimated using a Bayesian framework implemented using the R and Stan statistical computing platforms. See details in Supplementary Information, but the basic logic of the likelihood function can be understood, in its main principles, through these steps: 1) compute the time series *λ_t_* in a location using local climate time series and the growth-environment function, eq. 5 (or its multivariate generalization, Supplementary Information); 2) calculate the ltsgr in a location using eq. 4 and the *λ_t_*; and 3) compute the suitability score in the location using the ltsgr and eq. 6. The overall likelihood is the product of these suitability scores across all detection locations, times the product of 1 minus these scores across all (pseudo)absences for the species. We used uninformative priors for all parameters (Supplementary Information). Outputs of fitting include samples from the posterior distribution of the parameters *θ_3_*.

### Habitat suitability maps and species distributions

Given parameter estimates, 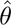*_3_*, a habitat suitability map for those parameters could be generated by carrying out the following steps for all locations across the region of interest: 1) compute the time series *λ_t_* in the location using local climate time series and the growth-environment function (eq. 5 or its multivariate generalization, Supplementary Information), with parameters taken from 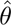*_3_*; 2) calculate the ltsgr in the location using eq. 4 and the *λ_t_*; 3) compute the habitat suitability, using the ltsgr and eq. 6, with parameters taken from 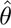*_3_*. For our simulation studies (see below), we used parameter point estimates for 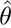*_3_*. For empirical species, instead, we computed a habitat suitability map for each of 300 posterior samples and averaged those maps; though results obtained in that way differed very little from simply using point estimates. XSDM therefore provides a suitability score for each location in geographic space, incorporating the influence of inter-annual climate variability. Thresholds can be applied to the habitat suitability map to provide a species geographic distributions/range map; we used the threshold that maximized the sum of sensitivity and specificity [47]. We emphasize that maps created in these ways show only projected suitability of each location with respect to environmental variables included in the model, ignoring possible dispersal barriers and range limitations which may be imposed by other species with which the focal species interacts. In other words, the maps represent the potential distribution of the species, given climatic conditions [4]. See Supplementary Information for additional details.

### Simulation studies

To assess whether XSDM could be used to accurately infer parameter values, we simulated data for virtual species using XSDM itself as the data-generating model and then fitted XSDM to these datasets. This is needed, in part, to make sure that the parameters of XSDM are “practically identifiable” [46]. We simulated climate time series of 40 years, comparable with available climate data, and explored four different scenarios differing in average climate and in its spatial heterogeneity. For each scenario, we ran 1,000 simulations, varying the total number of species presence records. XSDM performed similarly in the four scenarios, so we reported only one scenario in the main text. See Supplementary Information for additional details.

We ran an additional set of simulations to compare the performance of XSDM against traditional SDM approaches, namely: generalized additive models, maximum entropy, generalized linear models, and random forests. To avoid biasing these comparisons in favor of XSDM, we used a data generating model for these simulations which differed from the XSDM model. Specifically, the populations of the virtual species were modeled by

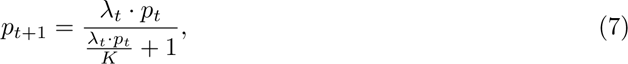

where *p* is the number of individuals, *K* is the carrying capacity of the population (*K* ∼ U(100, 200)) and *p_0_* ∼ U(5, 15). We calculated the habitat suitability score again using an inverse-logit function as in eq. 6 (with *b* = 1, *c* = 0, and *p_d_* = 1), but inputting the average simulated population density across the time period instead of the ltsgr. The suitability score was then used in a Bernoulli process to generate presences and absences, which were used as input for all SDMs. We compared the predictive performance of SDMs by plotting the predicted suitability against the true suitability. See previous section and Supplementary Information for additional details.

### XSDM for 10 species in GBIF

We used XSDM to estimate suitability maps and potential distributions of 10 species, chosen as illustrative examples from checklists of North American species (Supplementary Information for details). GBIF data were filtered to include only occurrences recorded after 1979 and with a spatial uncertainty ≤ 5 km. The remaining occurrence records were additionally filtered using the R package *CoordinateCleaner* [48] to remove spurious records. We sampled pseudo-absences up to 100 km distant from the minimum convex polygon containing all filtered presence records, assumed to be the maximum dispersal range of the species. We kept at most one presence and pseudo-absence record per grid cell. The number of pseudo-absences was set to three times the number of presences, i.e. we fixed the prevalence of all species at 0.25. These choices are fairly standard [49, 50].

As climatic predictors, we used detrended annual time series of bioclimatic variables related to temperature and precipitation, derived from CHELSA [51]: average annual temperature (BIO01); mean temperature of the warmest quarter (BIO10); mean temperature of the coldest quarter (BIO11); total annual precipitation (BIO12); precipitation of the wettest quarter (BIO16); and precipitation of the driest quarter (BIO17). For each species, we ran nine competing models using the multivariate version of XSDM and all combinations of one temperature and one precipitation variable. Models were removed from further consideration that did not meet criteria for adequate chain convergence in Stan, or which had inadequate effective sample size (all model fitting details in Supplementary Information). For each species, we then selected as the best model for that species the model that had the highest expected log-pointwise predictive density (ELPD) using leave-one-out cross-validation [52]. Species data are linked and some fitting results are tabulated in Supplementary Table 2. Posterior samples (see Supplementary Table 3 for their point-estimates) were extracted from the best model and used to project the distribution of the species in North America, both with and without the influence of climatic variability, as well as to calculate climate sensitivity of species (see below).

### Quantifying the influence of climate variability with Δ-XSDM

An important feature of XSDM is that it can directly quantify the influence of inter-annual climate variability on habitat suitability maps and species distribution maps. This is achieved by comparing the suitability maps or distributions described above with maps or distributions obtained in the same way but starting from artificial environmental time series which are constant in each location at the mean value for the location. The same posterior sample of *θ*^^^*_3_* was used, and, for the distribution, the same threshold was used. We call this procedure Δ-XSDM.

### Sensitivities

To assess and compare how species are likely to be affected by changes in average temperature versus by changes in the inter-annual variability of temperature, we calculated sensitivities of species’ geographic range sizes to such changes. Sensitivities were instantaneous rates of change (derivatives) of the area of species geographic ranges with respect to changes in the average or the standard deviation of the temperature time series influencing the species, expressed as a percentage of current range size (details in Supplementary Information). Averages and standard deviations of temperature time series both have units of *^◦^*C, so our sensitivity measures all had units of *^◦^*C*^−1^* and could be directly compared. Sensitivities were used instead of projections of range size change under specific scenarios of future altered climate to avoid uncertainties that come with such extrapolations.

### Software and data availability

Analyses were performed in the R v4.2.2 and Stan v2.33.0 programming languages for the simulations; and in R v4.4.2 and Stan v2.36.0 [53, 54] for the analyses of GBIF data, the latter at a resolution of 5 × 5 km*^2^*using Albers conic equal-area projection (EPSG:5070). Data and results that support the findings of this study are available on Zenodo https://zenodo.org/doi/10.5281/zenodo.13378593.

## 9 Acknowledgments

EB and BR acknowledge the support of the German Centre for Integrative Biodiversity Research (iDiv) Halle-Jena-Leipzig, funded by the German Research Foundation (DFG, FZT 118, 202548816). DCR was partly supported by The Alexander von Humboldt and James S McDonnell Foundations, the California Department of Fish and Wildlife, US National Science Foundation grants DEB-PCE 2414418 and BIO-OCE 2023474, and the University of Kansas.

## Author contribution

EB and DCR conceived of the idea and planned the analyses. EB, BR, AR, and DCR performed the analyses. EB and DCR wrote the first draft of the manuscript. All authors contributed to the final version of the manuscript.

