## Extended Data for "The impacts of climate variability on the niche concept and distributions of species"

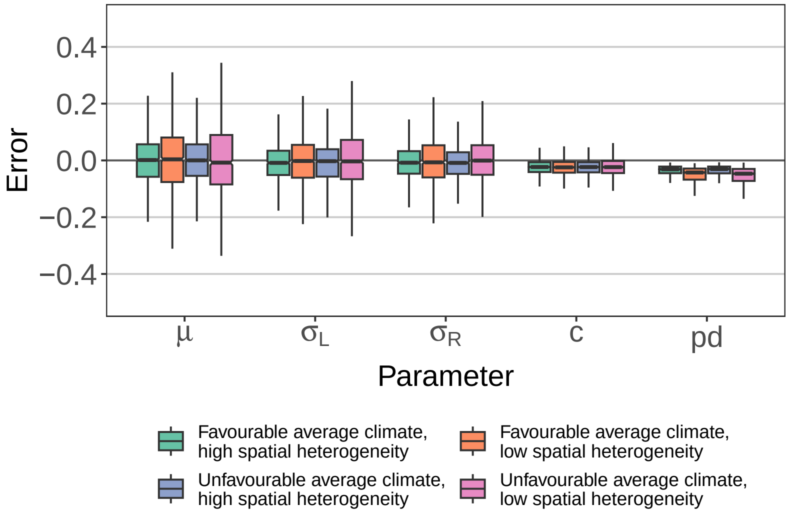


Figure 1: Distribution of errors in parameter values estimated by XSDM fitting for the first set of simulations we did. See Methods and the Supplementary informatio section "Simulations to assess whether XSDM ..." for details. The y-axis shows the distribution of the differences between the true and estimated parameter values across 1000 simulations. The central tendency of each distribution illustrates bias and the boxes and whiskers illustrate variance of the estimators. Colors show the four macro-climatic scenarios investigated (Supplementary information). Though only the dimension-reduced parameters $\theta_3$ could be estimated (see Methods), estimates of the parameters listed on the horizontal axis could be recovered using the known (true) values of $b$ and $\lambda_{\max}$.

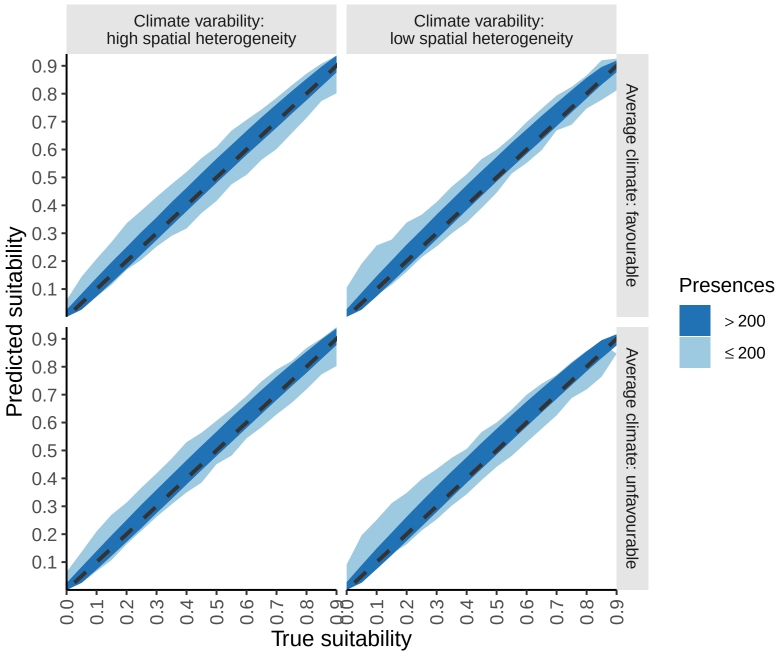


Figure 2: Results from the first set of simulations, similar to Fig. 3A in the main text, but now for all the macro-climatic scenarios. See the Supplementary information section "Simulations to assess whether XSDM . . . " for details.

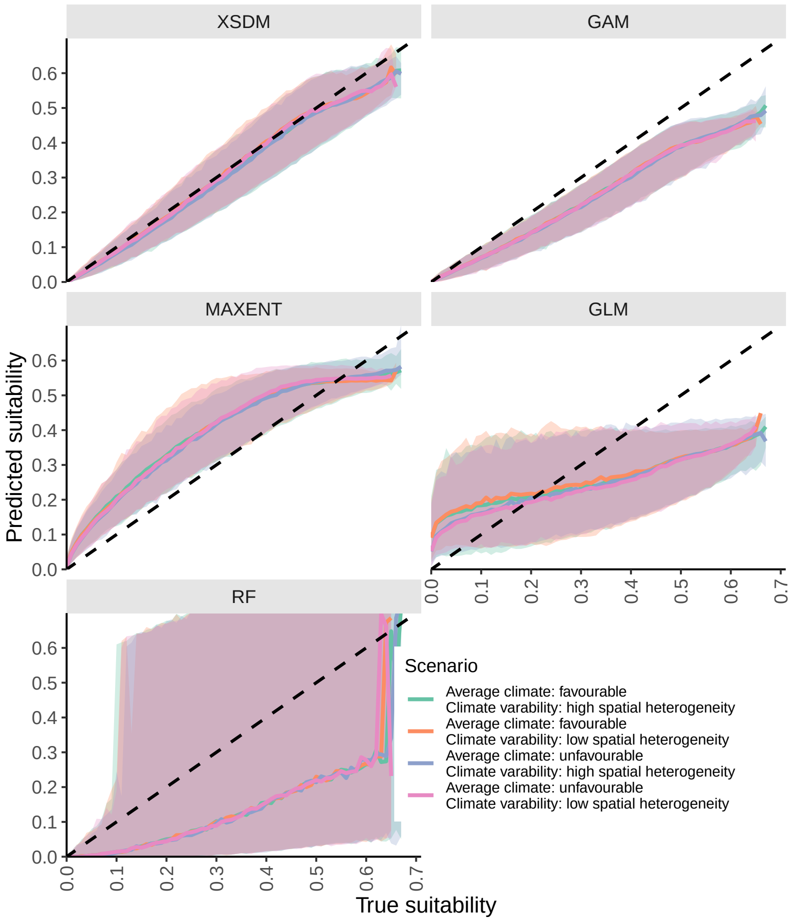


Figure 3: Results from the second set of simulations, similar to Fig. 3C in the main text, but now for all the macro-climatic scenarios. See the Supplementary information section "Simulations to benchmark XSDM..." for details.

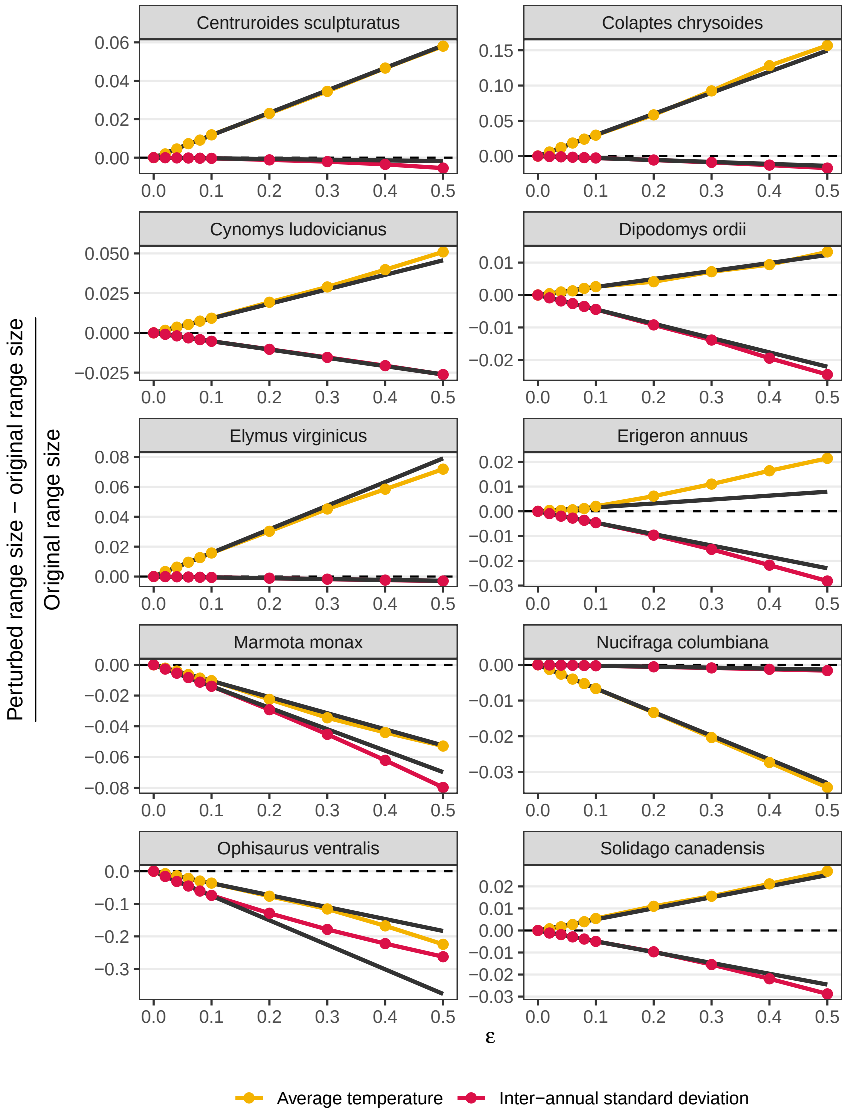


Figure 4: Simulated change in relative range size for a small increase of temperature average or inter-annual standard deviation. Circles and colored lines show the relative change for a small increase ($\epsilon$) of average temperature or inter-annual standard deviation; and the black solid line is the best-fit line, constrained to cross the origin, through points with $\epsilon \leq 0.1$. See Methods for details.

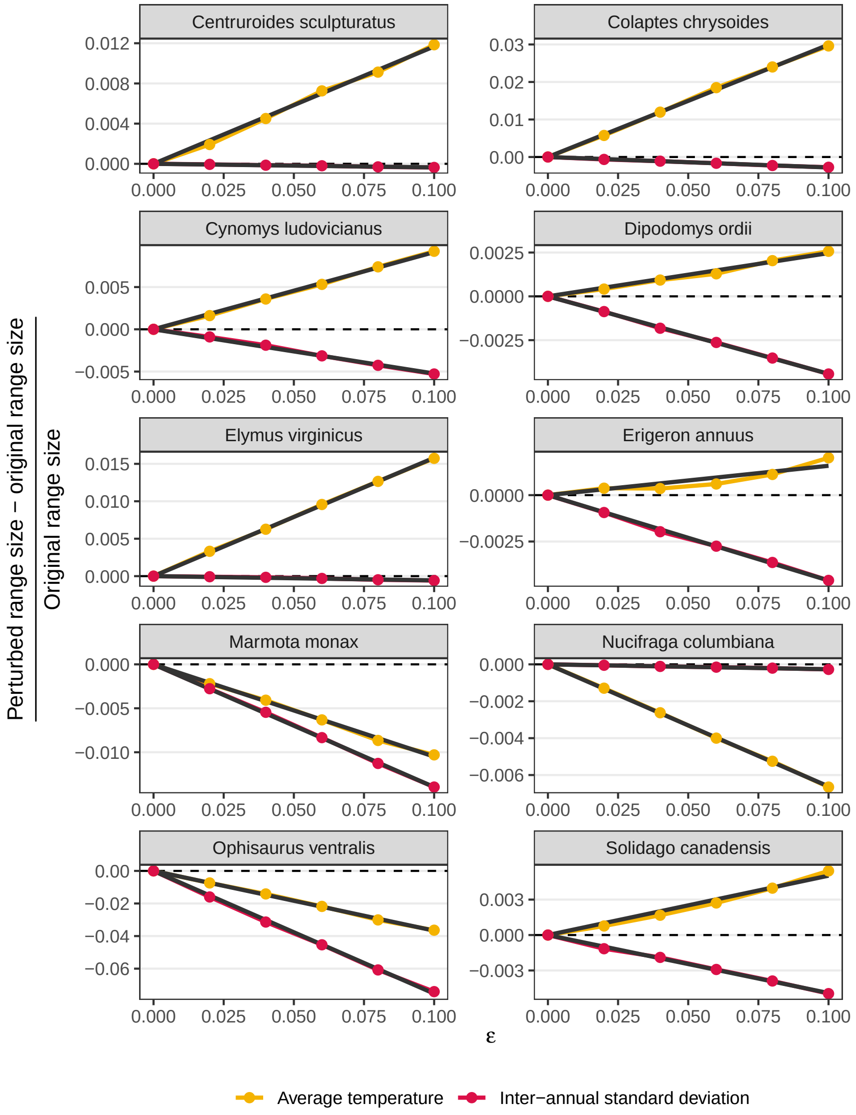


Figure 5: Simulated change in relative range size for a small increase of temperature average or inter-annual standard deviation for $\epsilon \leq 0.10$. Circles and colored lines show the relative change for a small increase ($\epsilon$) of average temperature or inter-annual standard deviation and the black solid line is the best-fit line crossing the origin. The trend between $\epsilon$ and the relative range change is approximately linear for all species, except for the increase of average temperature for *Erigeron annuus*. See Methods and Supplementary information for details.
