## Supplementary Information for "The impacts of climate variability on the niche concept and distributions of species"

### Contents

**1 Supplementary note: Fitting univariate XSDMs** 2

**2 Supplementary note: Fitting multivariate XSDMs** 3

**3 Supplementary note: Habitat suitability maps and species distributions** 6

**4 Supplementary note: Simulations to assess whether XSDM can be used to accurately infer model parameters** 6

**5 Supplementary note: Simulation to benchmark XSDM against other SDMs** 8

**6 Supplementary note: Species selected for illustrating XSDM, and data preparation** 9

**7 Supplementary note: Sensitivities** 11

**8 Supplementary figures** **12**

**9 Supplementary tables** **14**

### 1 Supplementary note: Fitting univariate XSDMs

We here describe the process of fitting the XSDM model to species occurrence data in the case where the growth-environment function, $\lambda_t$, is assumed to depend on only one environmental variable, $e_t$ (Methods). For typical applications, $\lambda_t$ will depend on multiple environmental variables, but describing the univariate case first allows for simpler exposition of main ideas.

Under the XSDM framework with univariate growth-environment function (Methods), the habitat suitability, for the focal species, at a location is

$$P(X = 1|\text{ltsgr}, b, c, p_d) = p_d \text{logit}^{-1}(b(\text{ltsgr} - c)) = \frac{p_d}{1 + \exp\left(-b(\text{ltsgr} - c)\right)}. \quad (1)$$

We then have

$$\text{ltsgr} = \overline{\log(\lambda_t)} \quad (2)$$

$$= \log(\lambda_{\max}) - \frac{1}{2}\overline{\left(\frac{e_t - \mu}{\sigma(e_t, \mu)}\right)^2}, \quad (3)$$

and therefore

$$P(X = 1|\text{ltsgr}, b, c, p_d) = P(X = 1|e_t, \lambda_{\max}, \mu, \sigma_L, \sigma_R, b, c, p_d) \quad (4)$$

$$= \frac{p_d}{1 + \exp\left(-b\left[\log(\lambda_{\max}) - \frac{1}{2}\overline{\left(\frac{e_t-\mu}{\sigma(e_t,\mu)}\right)^2} - c\right]\right)} \quad (5)$$

$$= \frac{p_d}{1 + \exp\left(\frac{b}{2}\overline{\left(\frac{e_t-\mu}{\sigma(e_t,\mu)}\right)^2} + b(c - \log(\lambda_{\max}))\right)} \quad (6)$$

$$= \frac{p_d}{1 + \exp\left(\frac{1}{2}\overline{\left(\frac{e_t-\mu}{\sigma(e_t,\mu)/\sqrt{b}}\right)^2} + b(c - \log(\lambda_{\max}))\right)}. \quad (7)$$

A likelihood function is established as $\prod_j P(X_j = 1) \prod_j (1 - P(X_j = 1))$, where the first product is over all detection locations and the second is over all locations of absence or pseudo-absence.

But eq. 7 shows that XSDM, as currently parameterized, is a structurally non-identifiable model [1] because some parameters occur together in that equation in a manner which allows them to trade off exactly against each other. For instance, $c$ and $\lambda_{\max}$ are confounded parameters because they only occur in the expression $c - \log(\lambda_{\max})$; increasing $c$ while increasing $\log(\lambda_{\max})$ by the same amount leads to the same likelihood value, so that the likelihood function, as currently written, cannot have a single maximum. Additionally, $\sigma_L$, $\sigma_R$, $b$, and $c - \log(\lambda_{\max})$ are also confounded. We reduce the parameter space to the parameters $\theta_3 = \{\mu, \tilde{\sigma}_L = \sigma_L/\sqrt{b}, \tilde{\sigma}_R = \sigma_R/\sqrt{b}, \tilde{c} = b(c - \log(\lambda_{\max})), p_d\}$. Then,

$$P(X = 1|e_t, \mu, \tilde{\sigma}_L, \tilde{\sigma}_R, \tilde{c}, p_d) = \frac{p_d}{1 + \exp\left[\frac{1}{2}\left(\frac{e_t - \mu}{\tilde{\sigma}(e_t, \mu)}\right)^2 + \tilde{c}\right]}, \tag{8}$$

where $\tilde{\sigma}(e_t, \mu) = \tilde{\sigma}_L$ for $e_t \leq \mu$ and $\tilde{\sigma}(e_t, \mu) = \tilde{\sigma}_R$ for $e_t > \mu$. This model can be fitted using occurrence data. It is not structurally non-identifiable, nor was it practically non-identifiable [1] for our data. See "Species selected for illustrating..." below, for information on priors.

### 2 Supplementary note: Fitting multivariate XSDMs

We now describe the process of fitting the XSDM model to species occurrence data when the growth-environment function, $\lambda_t$, is assumed to depend on $n > 1$ environmental variables, the components of an $n$-vector $\vec{e}_t$. The dynamical model and suitability-link function are the same as in Methods.

We start by defining the multivariate growth-environment function. Given an $n$-vector parameter $\vec{\mu} = (\mu_1, \ldots, \mu_n)$; two more $n$-vector parameters $\vec{\sigma}_L = (\sigma_{L,1}, \ldots, \sigma_{L,n})$ and $\vec{\sigma}_R = (\sigma_{R,1}, \ldots, \sigma_{R,n})$ with positive entries; and an $n \times n$ Cholesky factor of a correlation matrix, $L$, we define

$$\lambda_t = \lambda_{\max} \exp\left(-\frac{1}{2}\left[L^{-1}F_{\vec{\sigma}_L,\vec{\sigma}_R}(\vec{e}_t - \vec{\mu})\right]^t \left[L^{-1}F_{\vec{\sigma}_L,\vec{\sigma}_R}(\vec{e}_t - \vec{\mu})\right]\right), \tag{9}$$

where $F_{\vec{\sigma}_L,\vec{\sigma}_R}(\vec{u})_i = u_i/\sigma_{L,i}$ for $u_i < 0$ and $F_{\vec{\sigma}_L,\vec{\sigma}_R}(\vec{u})_i = u_i/\sigma_{R,i}$ for $u_i \geq 0$; and the superscript $t$ denotes matrix transpose. In the case of $n = 1$, the only possibility for $L$ is $L = 1$, and eq. (9) reduces to eq. 5 in Methods. At the end of this section we return to further justify eq. (9). Examples of $\lambda_t$ for the case $n = 2$ are in Fig. S2.

We then have

$$\begin{aligned} \text{ltsgr} &= \overline{\log(\lambda_t)} & (10) \\ &= \log(\lambda_{\max}) - \frac{1}{2}\overline{[L^{-1}F_{\vec{\sigma}_L,\vec{\sigma}_R}(\vec{e}_t - \vec{\mu})]^t \, [L^{-1}F_{\vec{\sigma}_L,\vec{\sigma}_R}(\vec{e}_t - \vec{\mu})]}, & (11) \end{aligned}$$

where the mean is, as usual, computed over time. Plugging this into the suitability-link function gives

$$
\begin{aligned}
P(X=1|\text{ltsgr},b,c,p_d) &= P(X=1|\vec{e}_t,\lambda_{\max},\vec{\mu},\vec{\sigma}_L,\vec{\sigma}_R,b,c,p_d) && (12)\\
&= \frac{p_d}{1+\exp\left(-b\left[\log(\lambda_{\max})-\frac{1}{2}\overline{[L^{-1}F_{\vec{\sigma}_L,\vec{\sigma}_R}(\vec{e}_t-\vec{\mu})]^t\,[L^{-1}F_{\vec{\sigma}_L,\vec{\sigma}_R}(\vec{e}_t-\vec{\mu})]}-c\right]\right)} && (13)\\
&= \frac{p_d}{1+\exp\left(\frac{b}{2}\overline{[L^{-1}F_{\vec{\sigma}_L,\vec{\sigma}_R}(\vec{e}_t-\vec{\mu})]^t\,[L^{-1}F_{\vec{\sigma}_L,\vec{\sigma}_R}(\vec{e}_t-\vec{\mu})]}+b(c-\log(\lambda_{\max}))\right)} && (14)\\
&= \frac{p_d}{1+\exp\left(\frac{1}{2}\overline{\left[L^{-1}\sqrt{b}F_{\vec{\sigma}_L,\vec{\sigma}_R}(\vec{e}_t-\vec{\mu})\right]^t\left[L^{-1}\sqrt{b}F_{\vec{\sigma}_L,\vec{\sigma}_R}(\vec{e}_t-\vec{\mu})\right]}+b(c-\log(\lambda_{\max}))\right)}. && (15)
\end{aligned}
$$

However,

$$
\begin{aligned}
\sqrt{b}F_{\vec{\sigma}_L,\vec{\sigma}_R}(\vec{u})_i &= \begin{cases}\sqrt{b}\frac{u_i}{\sigma_{L,i}} & \text{if } u_i<0\\ \sqrt{b}\frac{u_i}{\sigma_{R,i}} & \text{if } u_i\geq 0\end{cases} && (16)\\
&= \begin{cases}\frac{u_i}{\sigma_{L,i}/\sqrt{b}} & \text{if } u_i<0\\ \frac{u_i}{\sigma_{R,i}/\sqrt{b}} & \text{if } u_i\geq 0\end{cases} && (17)\\
&= F_{\vec{\sigma}_L/\sqrt{b},\vec{\sigma}_R/\sqrt{b}}(\vec{u})_i, && (18)
\end{aligned}
$$

so we have

$$
P(X=1|\text{ltsgr},b,c,p_d) = \frac{p_d}{1+\exp\left(\frac{1}{2}\overline{\left[L^{-1}F_{\vec{\sigma}_L/\sqrt{b},\vec{\sigma}_R/\sqrt{b}}(\vec{e}_t-\vec{\mu})\right]^t\left[L^{-1}F_{\vec{\sigma}_L/\sqrt{b},\vec{\sigma}_R/\sqrt{b}}(\vec{e}_t-\vec{\mu})\right]}+b(c-\log(\lambda_{\max}))\right)}. \quad (19)
$$

The likelihood function is again established as $\prod_j P(X_j=1)\prod_j(1-P(X_j=1))$, where the first product is over all detection locations and the second is over all locations of absence or pseudo-absence.

But eq. (19) shows that the model, as parameterized, is structurally non-identifiable. We reduce parameter space to the parameters $\theta_3=\{\vec{\mu},\tilde{\vec{\sigma}}_L=\vec{\sigma}_L/\sqrt{b},\tilde{\vec{\sigma}}_R=\vec{\sigma}_R/\sqrt{b},L,\tilde{c}=b(c-\log(\lambda_{\max})),p_d\}$.

Then,

$$P(X=1) = \frac{p_d}{1+\exp\left(\frac{1}{2}\left[L^{-1}F_{\tilde{\vec{\sigma}}_L,\tilde{\vec{\sigma}}_R}(\vec{e}_t-\vec{\mu})\right]^t\left[L^{-1}F_{\tilde{\vec{\sigma}}_L,\tilde{\vec{\sigma}}_R}(\vec{e}_t-\vec{\mu})\right]+\tilde{c}\right)}. \tag{20}$$

The resulting simplified model can be fitted using occurrence data. It is not structurally non-identifiable, nor was it practically non-identifiable for our data.

We now elaborate the reasoning behind the definition of the growth environment function described in eq. (9), above. In the univariate case (Methods and section 1 of this Supplementary information), it was convenient to describe the growth-environment function using an asymmetric generalization of a functional form also used for the probability density function (pdf) of a normal distribution,

$$\exp\left(-\frac{1}{2}\left(\frac{e_t-\mu}{\sigma}\right)^2\right). \tag{21}$$

Specifically,

$$\lambda_t = \lambda_{\max}\exp\left(-\frac{1}{2}\left(\frac{e_t-\mu}{\sigma(e_t,\mu)}\right)^2\right), \tag{22}$$

where $\sigma(e_t,\mu)=\sigma_L$ if $e_t \leq \mu$, and $\sigma(e_t,\mu)=\sigma_R$ if $e_t>\mu$. This functional form was convenient because annual net growth rate should decline monotonically to zero for extreme values of $e_t$, though it may do so asymmetrically. For similar reasons, the multivariate growth-environment function is an asymmetric generalization of a functional form also used in the pdf of the multivariate normal distribution,

$$\exp\left(-\frac{1}{2}(\vec{e}_t-\vec{\mu})^t\Sigma^{-1}(\vec{e}_t-\vec{\mu})\right) \tag{23}$$

where $\Sigma$ is a positive-definite covariance matrix. To define an asymmetric generalization of eq. (23), we first write $\Sigma = DCD$, where $C$ is a correlation matrix and $D$ is the diagonal matrix obtained by taking square roots of the diagonal entries of $\Sigma$. We then take the Cholesky decomposition $C = L_C L_C^t$, so that $\Sigma = DL_CL_C^tD^t$, and then

$$(\vec{e}_t-\vec{\mu})^t\Sigma^{-1}(\vec{e}_t-\vec{\mu}) = (\vec{e}_t-\vec{\mu})^t(DL_CL_C^tD^t)^{-1}(\vec{e}_t-\vec{\mu}) \tag{24}$$

$$= [L_C^{-1}D^{-1}(\vec{e}_t-\vec{\mu})]^t[L_C^{-1}D^{-1}(\vec{e}_t-\vec{\mu})]. \tag{25}$$

We note that

$$D^{-1}(\vec{e}_t-\vec{\mu}) = F_{\vec{\sigma},\vec{\sigma}}(\vec{e}_t-\vec{\mu}) \tag{26}$$

for $\vec{\sigma}$ equal to the diagonal of $D$. The asymmetric generalization simply replaces $F_{\vec{\sigma},\vec{\sigma}}(\vec{e}_t-\vec{\mu})$ by $F_{\vec{\sigma}_L,\vec{\sigma}_R}(\vec{e}_t-\vec{\mu})$.

### 3 Supplementary note: Habitat suitability maps and species distributions

Fitting XSDM with species occurrence data provides samples from the posterior distribution of $\theta_3$. We based habitat suitability and species distributions on those samples or on the parameter point estimate $\hat{\theta}_3$. For simulations, plugging point estimates, $\hat{\theta}_3$, into eq. 8 (for the univariate case) or eq. 20 (for the multivariate case) gave a function of the local environment time series that provided a habitat suitability score for every simulated location. For empirical species occurrence records, plugging 300 samples from the posterior distribution of $\theta_3$ into eq. 8 or 20 provided 300 functions of the local environment time series; for each location, these were averaged to produce the suitability map that was used. Species distribution maps can be constructed by thresholding this habitat suitability.

### 4 Supplementary note: Simulations to assess whether XSDM can be used to accurately infer model parameters

To assess whether XSDM can be used to accurately infer model parameters, we performed a simulation study where we repeatedly generated data for virtual species using the XSDM model, and then fitted XSDM to those simulated data. We used the XSDM model for both data generation and fitting because our aim, for these simulations, was to assess aspects of bias, variance and consistency of the parameter estimators provided by XSDM; but see below for additional simulations where the generating model differed from the XSDM model. The univariate version of the XSDM growth-environment function was used.

For each simulation, we first determined the growth-environment function of the virtual species for that simulation by sampling $\mu \sim \mathrm{U}(-0.5, 0.5)$, $\sigma_L \sim \mathrm{U}(0.5, 2)$, and $\sigma_R \sim \mathrm{U}(0.5, 2)$, where U represents the uniform distribution. The suitability-link parameters $b = 5$, $c = -0.5$, and $p_d = 1$, as well as $\lambda_{\max} = 1$, were used for all simulations in this section. The number of potential habitat patches, $J$, to be considered for an individual simulation was generated using $J \sim \mathrm{U}(10^2, 10^4)$.

We then created annual climatic time series for the $J$ locations based on four macro-climatic scenarios. To describe this process we first define some notation. Given an overall central-tendency parameter, $E$, and a spread parameter, $V$, location-specific central-tendency and spread parameters $E_j$ and $V_j$ were generated as $E_j \sim \mathrm{N}(E, 1.5^2)$ and $V_j \sim \mathrm{TN}(1, V^2)$ for $j = 1, \ldots, J$. Here, N represents the normal distribution, with mean and variance parameters, and TN is a normal distribution truncated at 0. Then, a length-40 time series $e_t^{(j)}$ was generated for each location, $j$, using 40 independent draws from $\mathrm{N}(E_j, V_j)$. The four macro-climatic scenarios were: A) environment was generally favorable ($E = 0$), and its temporal variability was relatively spatially homogeneous ($V = 5 \cdot 10^{-3}$); B) environment was favorable ($E = 0$) and its temporal variability was relatively spatially heterogeneous ($V = 0.5$); C) environment was generally unfavorable ($E = 1.5$) and its temporal variability was spatially homogeneous ($V = 5 \cdot 10^{-3}$); D) environment was unfavorable ($E = 1.5$) and its temporal variability was spatially heterogeneous ($V = 0.5$). For each macro-climatic scenario, 1000 simulations were carried out.

For each simulation, we calculated the annual net growth time series $\lambda_t$ and the ltsgr, and then obtained the habitat suitability of the species (eqs. 5, 4, and 6, respectively, of Methods). We used these suitability scores to sample from a Bernoulli distribution to obtain presences and absences which were the simulated data to which XSDM was subsequently fit.

For each simulated data set, we assessed if the XSDM fitting procedure converged, and the accuracy and precision which which parameters were recovered. Fit convergence was assessed using the Gelman-Rubin statistic $\hat{R}$, quantifying if the chains mixed well, and the bulk effective sample size ($n_{\mathrm{eff}}$), quantifying the sampling efficiency of the posteriors; fits that had $\hat{R} < 1.05$ and $n_{\mathrm{eff}} > 100$ were considered to have converged [2]. All of our 4,000 simulations converged. Model parameters, $\theta_3$, were fairly accurately recovered (Extended Data Fig. 1), with bias quite limited for the parameters more closely associated with the growth-environment function, which may be considered more important than parameters more closely associated with the suitability-link.

More importantly for our purposes, the accuracy with which fitted XSDM recovered habitat suitability maps was assessed, and was very good. True parameter values, $\theta_3$, were plugged into eq. 8, together with local environmental time series, to obtain true habitat suitability values for all locations. Estimated parameters, $\hat{\theta}_3$, were used in the same way to additionally determine habitat suitability as estimated from data by XSDM. We plotted estimated versus true suitability across all locations and simulations within a single macro-climatic scenario, separating plots into categories of numbers of species detections in the simulated data (Fig. 3A in the main text for one macro-climatic scenario, Extended Data Fig. 2 for all of them). We also plotted the difference between true and estimated habitat suitability scores as a function of number of species occurrences (Fig. 3B in the main text), finding, as expected, that accuracy improves with more occurrences, but already tends to be quite good with 200-300 occurrences. Results from the four macro-climates were quite similar (Extended Data Fig. 2), so in the main text we focused on only one of those scenarios.

### 5 Supplementary note: Simulation to benchmark XSDM against other SDMs

We performed an additional simulation study to assess how well XSDM performs compared to commonly used SDM approaches, including: generalized additive models (GAM), maximum entropy (Maxent), generalized linear models (GLM), and random forests (RF). Since statistical model miss-specification will commonly or always be a challenge faced by large-scale efforts to understand population dynamics, this simulation study used a data-generating model which differed from the XSDM model. It included aspects of density dependence, and species detection probability in a location was assumed to depend on density.

Specifically, we used $\lambda_t$ (eq. 5 in the main text) to calculate the annual population density ($p_t$) following

$$p_{t+1} = \frac{\lambda_t \cdot p_t}{\frac{\lambda_t \cdot p_t}{K} + 1}, \tag{27}$$

where $K$ is the carrying capacity of the population (drawn independently from $\mathrm{U}(100, 200)$ for each simulation) and $p_0$ is the initial condition (drawn independently from $\mathrm{U}(5, 15)$ for each simulation). We then calculated the probability of detection of the species in a location by plugging the average population density across the simulation time period (40 years) into the suitability-link function, in place of the ltsgr. The parameters $b = 1$, $c = 0$, and $p_d = 1$ were used for the suitability-link function for all simulations. We then used the suitability scores to sample from a Bernoulli distribution to obtain presences and absences for the virtual species for all simulated locations. The number of simulated locations was $J \sim \mathrm{U}(10^2, 10^4)$, as for the previous simulation study. Climatic time series were generated following four macro-climatic scenarios in the same manner as the previous simulation study.

We then fitted univariate XSDM and the four traditional SDM approaches to the simulated data. For XSDM, fit convergence was again assessed using the Gelman-Rubin statistic $\hat{R}$, quantifying if the chains mixed well, and the bulk effective sample size ($n_{\text{eff}}$), quantifying the sampling efficiency of the posteriors; fits that had $\hat{R} < 1.05$ and $n_{\text{eff}} > 100$ were considered to have converged [2]. All of our 4,000 simulations converged. For the traditional SDMs, we allowed as predictors the average climate value, and also its variance, and their interactions; AIC was computed for each possible combination of these predictors. This was done to assess if traditional SDMs can quantify the importance of inter-annual climatic variability for species suitability [3]. For the traditional SDMs, we retained the best (lowest AIC) model per algorithm. RF, GAM, and Maxent were fitted with the R packages *randomForest* [4], *mgcv* [5], and *maxnet* [6], respectively, using the default model parameterizations. As for the previous simulation study, predicted habitat suitability was plotted against true habitat suitability as assessed from the data-generating model. Results did not differ substantially among the four macro-climatic scenarios we explored (Extended Data Fig. 3), so results for only one scenario were presented in the main text.

### 6 Supplementary note: Species selected for illustrating XSDM, and data preparation

We selected species from taxonomic checklists of North America, e.g. `https://en.wikipedia.org/wiki/List_of_mammals_of_the_United_States`, avoiding species that were broadly distributed across the whole continent, as they were likely not limited by climatic suitability. For birds, we selected only species that did not have different distributional ranges between winter and summer. We downloaded the data from GBIF for each species separately, specifying as starting year of the dataset 1979 and a coordinate uncertainty $\leq 5$ km. Presences were cleaned using the R package *CoordinateCleaner* [7] to remove dubious locations, and were thinned to retain only one record per pixel. We then sampled pseudo-absences keeping the prevalence of the final dataset equal to 0.25, i.e. the number of pseudo-absences was three times the number of presences, making sure to have only one pseudo-absence per pixel.

Our goal was to obtain 10 species for which the version of XSDM described in this paper successfully fitted GBIF data for some choice of environmental variables. Since XSDM using the Lewontin-Cohen model may be too simple for some species (see Discussion for possible future XSDM extensions), we assessed if XSDM converged and produced a good fit by examining: i) the $\hat{R}$, effective sample size $n_{\text{eff}}$ and effective sample size in the tails, and overall convergence of the sampling; and ii) the ability of the fitted XSDM to predict GBIF data and our generated pseudo-absences, evaluated using the true-skill statistic (TSS). Specifically, for each of the nine models which were fitted for each species, models were eliminated that had: 10 or more divergences in the sampling chains; $\hat{R}$ for any of the parameters above 1.05; or bulk or tail $n_{\text{eff}}$ less than 400 for any of the parameters. We also inspected traceplots and pairs plots for each model. Species were then eliminated for which: there were not at least three models remaining; or TSS for the best-fitting model (as judged by expected log pointwise predictive density using leave-one-out cross-validation; see Methods) was less than 0.45. We ran XSDM for a total of 22 species (Table S2); 10 remained after the filtering described above. $\Delta$-XSDM was carried out and sensitivities were computed (Methods) for those species

Failures of convergence and low TSS values for some species were expected, and could have happened because: i) the climate variables we used (Methods) did not explain the distribution of those species; ii) the Lewontin-Cohen model was an overly simplistic model of those species' demography; or iii) longer sampling chains were needed. Because our main research goal was to understand the impact of inter-annual climate variability on species distributions, for our purposes it was sufficient to identify 10 species for which the simple version of XSDM we have described was an adequate model. Studies seeking to understand the distributions of specific species should consider a wider selection of environmental variables and may need to consider (st)age structured demographic models (Discussion), both of which should be priorities of future work.

We specified uninformative priors for all parameters. For $\mu_i$ (the $i$th component of $\vec{\mu}$), the prior was a normal distribution with mean equal to the average of the set $E_i$ consisting of all values $e_{i,t}$ across all locations of presence of the species and across the years considered. Here, $e_{i,t}$ is the $i$th component of $\vec{e}_t$. The standard deviation parameter of the prior of $\mu_i$ was taken to be 10 times the standard deviation of the values in $E_i$. The prior for the $i$th component $\tilde{\sigma}_{L,i}$ of $\vec{\tilde{\sigma}}_L$ was an exponential distribution with rate parameter equal to the inverse of the standard deviation of the values in $E_i$; and likewise for the $i$th component $\tilde{\sigma}_{R,i}$ of $\vec{\tilde{\sigma}}_R$. The prior for $\tilde{c}$ was normal with mean 0 and standard deviation 10. The prior for $L$ was the Lewandowski-Kurowicka-Joe distribution with parameter 2. The prior for $p_d$ was uniform on the interval from 0 to 1.

#### 7 Supplementary note: Sensitivities

For a temperature time series $T_t$, we simulated a small increase in its average by adding $\epsilon$ to each annual value: $T_t^\epsilon = T_t + \epsilon$. We simulated a small increase in its inter-annual variability by multiplying $T_t - \overline{T_t}$ by $\delta$: $T_t^\delta = \left(T_t - \overline{T_t}\right)\delta + \overline{T_t}$. We chose $\delta = 1 + \epsilon\left(\overline{\sqrt{\mathrm{Var}(T_t)}}\right)^{-1}$, where $\overline{\sqrt{\mathrm{Var}(T_t)}}$ is the inter-annual standard deviation averaged across all locations, so that the increase in inter-annual temperature variability was comparable to that of its average. Specifically, for an increase of $\epsilon$ in average temperature, the inter-annual standard deviation averaged across all locations also increases by $\epsilon$:

$$\overline{\sqrt{\mathrm{Var}(T_t^\delta)}} = \overline{\sqrt{\mathrm{Var}(T_t)}} + \epsilon \Leftrightarrow \delta\overline{\sqrt{\mathrm{Var}(T_t)}} = \overline{\sqrt{\mathrm{Var}(T_t)}} + \epsilon \Leftrightarrow \delta = 1 + \epsilon\left(\overline{\sqrt{\mathrm{Var}(T_t)}}\right)^{-1}. \tag{28}$$

We then used the resulting time series $T_t^\epsilon$ and $T_t^\delta$ to calculate the habitat suitability and distribution of the species as described in Methods, separately for $\epsilon$ equal to 0.02, 0.04, 0.06, 0.08, 0.10, 0.20, 0.30, 0.40, 0.50 °C. We denote by $A_{\mathrm{avg},\epsilon}$ the area of the species distributional range under a change to the average temperature of $\epsilon$; and we denote by $A_{\mathrm{sd},\epsilon}$ the area of the species distributional range under a change to the standard deviation of temperature of $\epsilon$. Here, $A_{\mathrm{avg},0}$ and $A_{\mathrm{sd},0}$ both correspond to range size under current conditions, a quantity which we denote $A_0$.

Sensitivities were the derivatives of the quantities $\Delta R_{\mathrm{avg},\epsilon} = (A_{\mathrm{avg},\epsilon} - A_0)/A_0$ and $\Delta R_{\mathrm{sd},\epsilon} = (A_{\mathrm{sd},\epsilon} - A_0)/A_0$ evaluated at $\epsilon = 0$, and were approximated as follows based on the quantities computed as described in the previous paragraph. We plotted $\Delta R_{\mathrm{avg},\epsilon}$ and $\Delta R_{\mathrm{sd},\epsilon}$ against $\epsilon$ for the $\epsilon$ values listed above and for $\epsilon = 0$ (Extended Data Figs 4, 5). All plots except the plot of $\Delta R_{\mathrm{avg},\epsilon}$ for *Erigeon annuus* were close to linear for the values of $\epsilon$ in $[0, 0.1]$, so sensitivities were approximated as regression slopes of $\Delta R_{\mathrm{avg},\epsilon}$ or $\Delta R_{\mathrm{sd},\epsilon}$ against $\epsilon$ using those values of $\epsilon$. The regressions used were constrained to pass through $(0, 0)$. For the plot of $\Delta R_{\mathrm{avg},\epsilon}$ for *Erigeon annuus*, we approximated the sensitivity as $\Delta R_{\mathrm{avg},0.02}/0.02$.

### 8 Supplementary figures

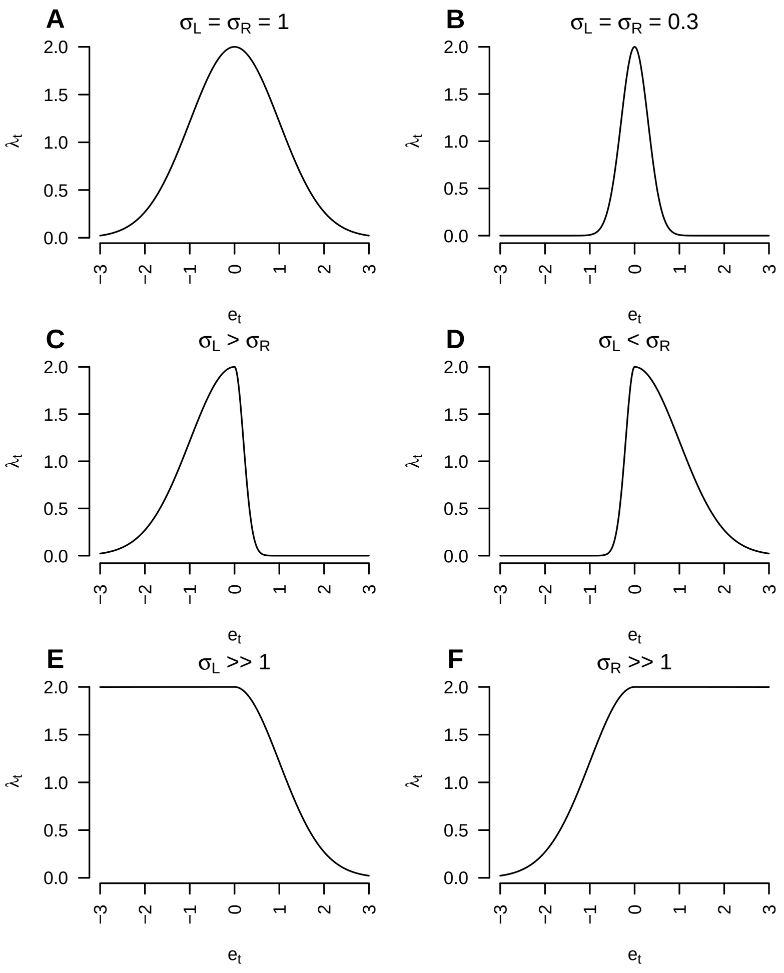


Figure S1: Examples of the possible shapes of the univariate version of the growth-environment function used in XSDM. **A**) A symmetric, broad response. **B**) A symmetric, narrow response. **C, D**) Asymmetric responses. **E**) A left-saturating response. **F**) A right-saturating response. Response optimum and maximum growth were always $\mu = 0$ and $\lambda_{max} = 2$, respectively for these examples.

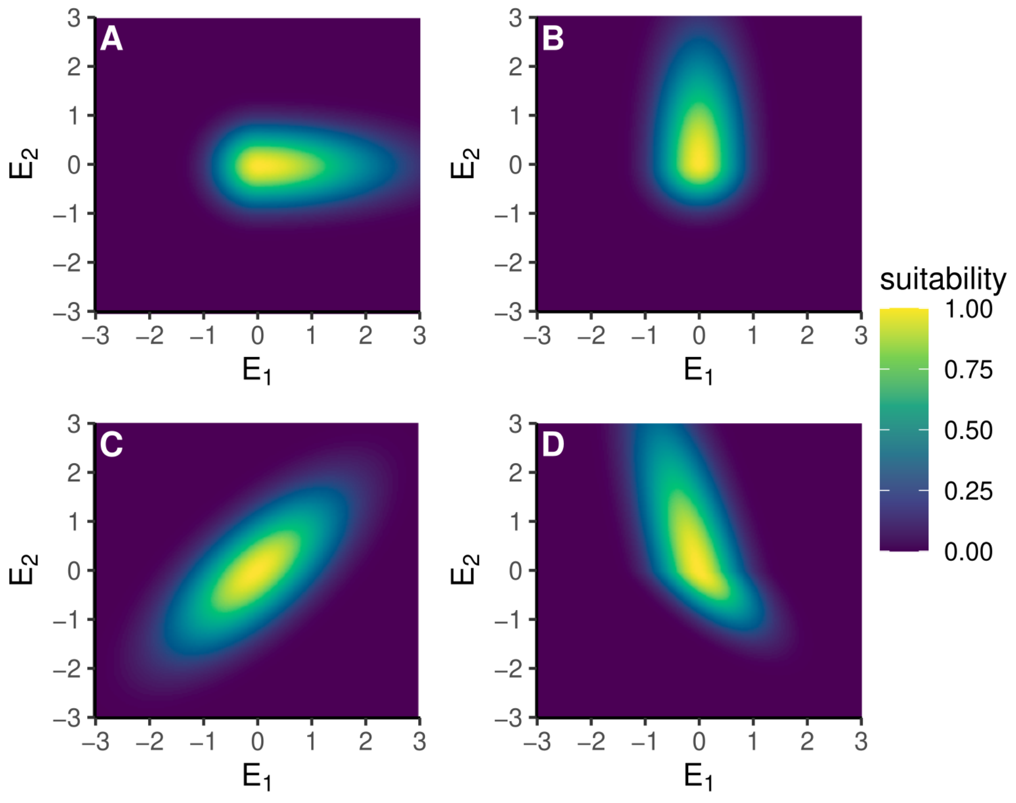


Figure S2: Examples of some of the possible shapes of multivariate growth-environment functions used in XSDM. All responses have optimal conditions $\mu = 0, 0$, but they differ in $\sigma_L$, $\sigma_R$, and $R = LL^t$. **A**) $\sigma_L = (0.5, 0.5)$, $\sigma_R = (1.5, 0.5)$, and $R_{12} = 0$. **B**) $\sigma_L = (0.5, 0.5)$, $\sigma_R = (0.5, 1.5)$, and $R_{12} = 0$. **C**) $\sigma_L = (1.0, 1.0)$, $\sigma_R = (1.0, 1.0)$, and $R_{12} = 0.7$. **D**) $\sigma_L = (0.7, 0.7)$, $\sigma_R = (0.7, 2.0)$, and $R_{12} = -0.7$.

### 9 Supplementary tables

Table S1: Sensitivity to a small increase of average temperature or its inter-annual standard deviation for the 10 species fitted using XSDM.

| **Species** | **increase in average** | **increase in inter-annual standard deviation** |
|---|---|---|
| *Centruroides sculpturatus* | 0.117 | -0.004 |
| *Colaptes chrysoides* | 0.299 | -0.028 |
| *Cynomys ludovicianus* | 0.091 | -0.052 |
| *Dipodomys ordii* | 0.025 | -0.044 |
| *Elymus virginicus* | 0.158 | -0.006 |
| *Erigeron annuus* | 0.019 | -0.047 |
| *Marmota monax* | -0.105 | -0.140 |
| *Nucifraga columbiana* | -0.066 | -0.003 |
| *Ophisaurus ventralis* | -0.367 | -0.753 |
| *Solidago canadensis* | 0.050 | -0.049 |

Table S2: Species modelled using XSDM and GBIF data. TSS was not computed for species for which fewer than three models converged, and the Δ-XSDM value was not computed unless convergence for at least three models occurred and TSS $\geq 0.45$ for the best model; see the section "Species selected for illustrating..." above.

| **Species** | **taxonomic class** | **TSS** | **doi** | **n GBIF** | **$\Delta-$XSDM** |
|---|---|---|---|---|---|
| *Ambystoma tigrinum* | Amphibia | 0.34 | `https://doi.org/10.15468/dl.w52dqh` | 481 | - |
| *Amphiuma means* | Amphibia | 0.34 | `https://doi.org/10.15468/dl.67b4ks` | 262 | - |
| *Blarina carolinensis* | Mammalia | 0.39 | `https://doi.org/10.15468/dl.k65m4y` | 289 | - |
| *Centruroides sculpturatus* | Arachnida | 0.56 | `https://doi.org/10.15468/dl.85d6g7` | 346 | 0.347 |
| *Colaptes chrysoides* | Aves | 0.65 | `https://doi.org/10.15468/dl.burbeb` | 299 | 0.316 |
| *Cryptotis parva* | Mammalia | 0.36 | `https://doi.org/10.15468/dl.7yybv7` | 375 | - |
| *Cynomys ludovicianus* | Mammalia | 0.51 | `https://doi.org/10.15468/dl.pekxx2` | 703 | 0.363 |
| *Dipodomys ordii* | Mammalia | 0.46 | `https://doi.org/10.15468/dl.mxndbx` | 627 | 0.209 |
| *Elymus virginicus* | Liliopsida | 0.52 | `https://doi.org/10.15468/dl.ztdqbh` | 1799 | 0.090 |
| *Erigeron annuus* | Magnoliopsida | 0.51 | `https://doi.org/10.15468/dl.ztdqbh` | 2619 | 0.132 |
| *Heloderma suspectum* | Squamata | - | `https://doi.org/10.15468/dl.8p7qks` | 77 | - |
| *Lampropeltis calligaster* | Squamata | 0.33 | `https://doi.org/10.15468/dl.h8ez2c` | 2088 | - |
| *Marmota monax* | Mammalia | 0.52 | `https://doi.org/10.15468/dl.szqtnq` | 5457 | 0.184 |
| *Nucifraga columbiana* | Aves | 0.49 | `https://doi.org/10.15468/dl.jr5s7f` | 1346 | 0.010 |
| *Ophisaurus ventralis* | Squamata | 0.54 | `https://doi.org/10.15468/dl.4ghjbe` | 682 | 0.447 |
| *Photinus pyralis* | Insecta | 0.41 | `https://doi.org/10.15468/dl.mrnjua` | 1919 | - |
| *Phrynosoma cornutum* | Squamata | 0.41 | `https://doi.org/10.15468/dl.dy3ky5` | 753 | - |
| *Plestiodon obsoletus* | Squamata | 0.36 | `https://doi.org/10.15468/dl.b5ckxs` | 459 | - |
| *Sauromalus ater* | Squamata | - | `https://doi.org/10.15468/dl.yedchy` | 468 | - |
| *Sceloporus poinsettii* | Squamata | 0.29 | `https://doi.org/10.15468/dl.hu6q5c` | 488 | - |
| *Solidago canadensis* | Magnoliopsida | 0.60 | `https://doi.org/10.15468/dl.asjwpe` | 2153 | 0.121 |
| *Sylvilagus nuttallii* | Mammalia | 0.31 | `https://doi.org/10.15468/dl.zzfu2r` | 650 | - |

Table S3: Best-model parameter point estimates of XSDM for the 10 species for which fitting converged for at least three models and TSS $\geq 0.45$ for the best model. The numbers in the subscripts refer to the climatic variables, with $_1$ indicating the temperature and $_2$ the precipitation variable. The best models had as predictors: The average temperature (BIO01) and the total precipitation (BIO12) for *Centruroides sculpturatus* and *Colaptes chrysoides*. BIO01 and the precipitation of the wettest quarter (BIO16) for *Dipodomys ordii*. BIO01 and the precipitation of the driest quarter (BIO17) for *Erigeron annuus* and *Solidago canadensis*. The temperature of the warmest quarter (BIO10) and BIO12 for *Elymus virginicus*. BIO10 and BIO16 for *Ophisaurus ventralis*. BIO10 and BIO17 for *Marmota monax* and *Nucifraga columbiana*. BIO11 and BIO16 for *Cynomys ludovicianus*. $L_{i,j}$ is the $i, j$ entry of the Cholesky factor ($L$ is constrained to have $L_{1,1} = 1$ and $L_{1,2} = 0$ because it is a Cholesky factor of a correlation matrix). Precipitation variables were divided by $10^4$ before fitting. Table values for parameters $\sigma$ refer to the corresponding reduced parameters $\tilde{\sigma}$, and table values for $c$ refer to the reduced parameter $\tilde{c}$.

| **Species** | **parameter** | **mean** | **sd** |
|---|---|---|---|
| *Centruroides sculpturatus* | $\mu_1$ | 20.482 | 0.815 |
| *Centruroides sculpturatus* | $\mu_2$ | 5.379 | 0.659 |
| *Centruroides sculpturatus* | $\sigma_{l,1}$ | 4.495 | 0.507 |
| *Centruroides sculpturatus* | $\sigma_{l,2}$ | 2.090 | 0.303 |
| *Centruroides sculpturatus* | $\sigma_{r,1}$ | 1.961 | 0.448 |
| *Centruroides sculpturatus* | $\sigma_{r,2}$ | 1.068 | 0.224 |
| *Centruroides sculpturatus* | $L_{2,1}$ | -0.648 | 0.059 |
| *Centruroides sculpturatus* | $L_{2,2}$ | 0.757 | 0.050 |
| *Centruroides sculpturatus* | $c$ | -1.617 | 0.334 |
| *Centruroides sculpturatus* | $p_d$ | 0.937 | 0.052 |
| *Colaptes chrysoides* | $\mu_1$ | 23.296 | 0.832 |
| *Colaptes chrysoides* | $\mu_2$ | 2.591 | 0.253 |
| *Colaptes chrysoides* | $\sigma_{l,1}$ | 3.317 | 0.381 |
| *Colaptes chrysoides* | $\sigma_{l,2}$ | 0.630 | 0.100 |
| *Colaptes chrysoides* | $\sigma_{r,1}$ | 2.677 | 1.803 |

| | | | |
|---|---|---|---|
| *Colaptes chrysoides* | $\sigma_{r,2}$ | 1.856 | 0.223 |
| *Colaptes chrysoides* | $L_{2,1}$ | -0.211 | 0.102 |
| *Colaptes chrysoides* | $L_{2,2}$ | 0.972 | 0.025 |
| *Colaptes chrysoides* | $c$ | -1.144 | 0.232 |
| *Colaptes chrysoides* | $p_d$ | 0.948 | 0.045 |
| *Cynomys ludovicianus* | $\mu_1$ | 5.302 | 0.963 |
| *Cynomys ludovicianus* | $\mu_2$ | 0.590 | 0.033 |
| *Cynomys ludovicianus* | $\sigma_{l,1}$ | 4.814 | 0.383 |
| *Cynomys ludovicianus* | $\sigma_{l,2}$ | 0.082 | 0.012 |
| *Cynomys ludovicianus* | $\sigma_{r,1}$ | 3.663 | 0.737 |
| *Cynomys ludovicianus* | $\sigma_{r,2}$ | 0.309 | 0.021 |
| *Cynomys ludovicianus* | $L_{2,1}$ | 0.119 | 0.052 |
| *Cynomys ludovicianus* | $L_{2,2}$ | 0.992 | 0.007 |
| *Cynomys ludovicianus* | $c$ | -1.826 | 0.435 |
| *Cynomys ludovicianus* | $p_d$ | 0.851 | 0.089 |
| *Dipodomys ordii* | $\mu_1$ | 13.923 | 0.678 |
| *Dipodomys ordii* | $\mu_2$ | 0.138 | 0.012 |
| *Dipodomys ordii* | $\sigma_{l,1}$ | 5.535 | 0.655 |
| *Dipodomys ordii* | $\sigma_{l,2}$ | 0.008 | 0.003 |
| *Dipodomys ordii* | $\sigma_{r,1}$ | 3.818 | 0.444 |
| *Dipodomys ordii* | $\sigma_{r,2}$ | 0.510 | 0.025 |
| *Dipodomys ordii* | $L_{2,1}$ | 0.196 | 0.111 |
| *Dipodomys ordii* | $L_{2,2}$ | 0.974 | 0.024 |
| *Dipodomys ordii* | $c$ | -0.479 | 0.136 |
| *Dipodomys ordii* | $p_d$ | 0.948 | 0.045 |
| *Elymus virginicus* | $\mu_1$ | 96.187 | 13.563 |
| *Elymus virginicus* | $\mu_2$ | 13.290 | 0.365 |
| *Elymus virginicus* | $\sigma_{l,1}$ | 17.349 | 1.660 |
| *Elymus virginicus* | $\sigma_{l,2}$ | 2.021 | 0.090 |
| *Elymus virginicus* | $\sigma_{r,1}$ | 3.610 | 3.568 |

| | | | |
|---|---|---|---|
| *Elymus virginicus* | $\sigma_{r,2}$ | 13.073 | 3.918 |
| *Elymus virginicus* | $L_{2,1}$ | 0.453 | 0.038 |
| *Elymus virginicus* | $L_{2,2}$ | 0.891 | 0.019 |
| *Elymus virginicus* | $c$ | -9.039 | 1.653 |
| *Elymus virginicus* | $p_d$ | 0.967 | 0.029 |
| *Erigeron annuus* | $\mu_1$ | 8.487 | 0.213 |
| *Erigeron annuus* | $\mu_2$ | 0.397 | 0.031 |
| *Erigeron annuus* | $\sigma_{l,1}$ | 1.949 | 0.114 |
| *Erigeron annuus* | $\sigma_{l,2}$ | 0.087 | 0.012 |
| *Erigeron annuus* | $\sigma_{r,1}$ | 4.319 | 0.192 |
| *Erigeron annuus* | $\sigma_{r,2}$ | 0.702 | 0.167 |
| *Erigeron annuus* | $L_{2,1}$ | 0.325 | 0.052 |
| *Erigeron annuus* | $L_{2,2}$ | 0.944 | 0.018 |
| *Erigeron annuus* | $c$ | -0.617 | 0.191 |
| *Erigeron annuus* | $p_d$ | 0.911 | 0.062 |
| *Marmota monax* | $\mu_1$ | 22.184 | 0.157 |
| *Marmota monax* | $\mu_2$ | 0.685 | 0.021 |
| *Marmota monax* | $\sigma_{l,1}$ | 2.412 | 0.082 |
| *Marmota monax* | $\sigma_{l,2}$ | 0.198 | 0.009 |
| *Marmota monax* | $\sigma_{r,1}$ | 2.083 | 0.094 |
| *Marmota monax* | $\sigma_{r,2}$ | 0.242 | 0.031 |
| *Marmota monax* | $L_{2,1}$ | 0.041 | 0.026 |
| *Marmota monax* | $L_{2,2}$ | 0.999 | 0.001 |
| *Marmota monax* | $c$ | -0.948 | 0.089 |
| *Marmota monax* | $p_d$ | 0.970 | 0.026 |
| *Nucifraga columbiana* | $\mu_1$ | -15.109 | 13.504 |
| *Nucifraga columbiana* | $\mu_2$ | -0.161 | 0.431 |
| *Nucifraga columbiana* | $\sigma_{l,1}$ | 3.688 | 3.618 |
| *Nucifraga columbiana* | $\sigma_{l,2}$ | 0.301 | 0.271 |
| *Nucifraga columbiana* | $\sigma_{r,1}$ | 8.499 | 1.563 |

| | | | |
|---|---|---|---|
| *Nucifraga columbiana* | $\sigma_{r,2}$ | 0.318 | 0.062 |
| *Nucifraga columbiana* | $L_{2,1}$ | 0.078 | 0.273 |
| *Nucifraga columbiana* | $L_{2,2}$ | 0.957 | 0.050 |
| *Nucifraga columbiana* | $c$ | -7.413 | 3.487 |
| *Nucifraga columbiana* | $p_d$ | 0.687 | 0.056 |
| *Ophisaurus ventralis* | $\mu_1$ | 28.974 | 0.988 |
| *Ophisaurus ventralis* | $\mu_2$ | 4.729 | 1.116 |
| *Ophisaurus ventralis* | $\sigma_{l,1}$ | 0.748 | 0.107 |
| *Ophisaurus ventralis* | $\sigma_{l,2}$ | 0.636 | 0.126 |
| *Ophisaurus ventralis* | $\sigma_{r,1}$ | 0.747 | 0.692 |
| *Ophisaurus ventralis* | $\sigma_{r,2}$ | 0.536 | 0.508 |
| *Ophisaurus ventralis* | $L_{2,1}$ | 0.719 | 0.111 |
| *Ophisaurus ventralis* | $L_{2,2}$ | 0.678 | 0.107 |
| *Ophisaurus ventralis* | $c$ | -12.692 | 4.038 |
| *Ophisaurus ventralis* | $p_d$ | 0.542 | 0.024 |
| *Solidago canadensis* | $\mu_1$ | 7.857 | 0.247 |
| *Solidago canadensis* | $\mu_2$ | 0.436 | 0.030 |
| *Solidago canadensis* | $\sigma_{l,1}$ | 2.718 | 0.126 |
| *Solidago canadensis* | $\sigma_{l,2}$ | 0.145 | 0.013 |
| *Solidago canadensis* | $\sigma_{r,1}$ | 4.207 | 0.167 |
| *Solidago canadensis* | $\sigma_{r,2}$ | 0.423 | 0.071 |
| *Solidago canadensis* | $L_{2,1}$ | 0.269 | 0.031 |
| *Solidago canadensis* | $L_{2,2}$ | 0.962 | 0.009 |
| *Solidago canadensis* | $c$ | -0.925 | 0.070 |
| *Solidago canadensis* | $p_d$ | 0.990 | 0.009 |

[5] S.N Wood. *Generalized Additive Models: An Introduction with R*. Chapman and Hall/CRC, 2 edition, 2017.

[6] Steven Phillips. *maxnet: Fitting 'Maxent' Species Distribution Models with 'glmnet'*, 2021. R package version 0.1.4.

[7] Alexander Zizka, Daniele Silvestro, Tobias Andermann, Josué Azevedo, Camila Duarte Ritter, Daniel Edler, Harith Farooq, Andrei Herdean, María Ariza, Ruud Scharn, et al. Coordinatecleaner: Standardized cleaning of occurrence records from biological collection databases. *Methods in Ecology and Evolution*, 10(5):744–751, 2019.
